# Integrating tracking and resight data from breeding Painted Bunting populations enables unbiased inferences about migratory connectivity and winter range survival

**DOI:** 10.1101/2020.07.23.217554

**Authors:** Clark S. Rushing, Aimee M. Van Tatenhove, Andrew Sharp, Viviana Ruiz-Gutierrez, Mary C. Freeman, Paul W. Sykes, Aaron M. Given, T. Scott Sillett

## Abstract

Archival geolocators have transformed the study of small, migratory organisms but analysis of data from these devices requires bias correction because tags are only recovered from individuals that survive and are re-captured at their tagging location. We show that integrating geolocator recovery data and mark-resight data enables unbiased estimates of both migratory connectivity between breeding and non-breeding populations and region-specific survival probabilities for wintering locations. Using simulations, we first demonstrate that an integrated Bayesian model returns unbiased estimates of transition probabilities between seasonal ranges and to determine how different sampling designs influence estimability of transition probabilities. We then parameterize the model with tracking data and mark-resight data from declining Painted Bunting (*Passerina ciris*) populations breeding in the eastern United States, hypothesized to be threatened by the illegal pet trade in parts of their Caribbean, non-breeding range. Consistent with this hypothesis, we found that male buntings wintering in Cuba were 20% less likely to return to the breeding grounds than birds wintering elsewhere in their range. Improving inferences from archival tags through proper data collection and further development of integrated models will advance our understanding of full annual cycle ecology of migratory species.

## Introduction

Rapid advances and miniaturization of tracking technology in recent decades have allowed us to quantify seasonal migrations of many terrestrial and aquatic species (Eckert & Stewart, 2001; Domeier & Nasby-Lucas, 2013; Jiménez López, Palacios, Jaramillo Legorreta, Urbán R., & Mate, 2019), to discover previously unknown migration routes (Sawyer, Kauffman, Nielson, & Horne, 2009; Smith et al., 2014; Naidoo et al., 2016), and to identify critical migration stopover and non-breeding areas (Richter & Cumming, 2008; Delmore, Fox, & Irwin, 2012; Cooper, Ewert, Jr, Helmer, & Marra, 2019). This information is essential for understanding population dynamics, disease transmission, range shifts, resource use, and management of vulnerable species or populations. Relatively large species (> 60g body mass) can be tracked with geolocation tags capable of transmitting location data to satellites (Scarpignato et al., 2016), but mapping seasonal movements and winter quarters for the great majority of migratory vertebrate species requires miniature archival geolocators (hereafter “geolocators”) that store location data internally and must be recovered from surviving individuals (Fraser et al., 2012; Hallworth & Marra, 2015; Peterson et al., 2015).

Although geolocators have been revolutionary for the study of small migratory organisms (McKinnon & Love, 2018), interpretation of migration patterns from the observed data must be done with care. In particular, because geolocators do not transmit data, observed migration data can only come from individuals that survive multiple migratory and stationary periods, return to their tagging location, and are recaptured. Individuals that do not survive at any stage of the annual cycle will not be represented. This form of *survivorship bias* is problematic for inferring migration patterns if certain migration routes have lower survival than others. For example, if the non-breeding range of a migratory species consists of two regions, one with high survival and one with low survival, individuals that migrate to the low-survival region will be less likely to return to their breeding site than individuals from the high-survival region. Hence, individuals from the low-survival region will be under-represented in the observed data relative their actual proportion. Estimates of transition probabilities (i.e., the probability that an individual from breeding site *i* migrates to non-breeding region *j*) will therefore be under-estimated for low-survival regions, and over-estimated for high-survival regions.

An analogous bias must be accounted for when estimating movement rates from band recoveries or re-sights with geographic variation in recovery/re-sight probabilities (Brownie, Hines, Nichols, Pollock, & Hestbeck, 1993; Nichols et al., 1995; Cohen, Hostetler, Royle, & Marra, 2014). Although a long history of model development is available to estimate these observation probabilities in dead-recovery and live-re-sight studies (reviewed by Korner-Nievergelt et al. (2010)), we lack an equivalent approach to account for the effects of survivorship bias in movement studies based on data from geolocators. Survivalship bias is an inevitable outcome of using archival geolocators when survival differs among migration routes and is therefore likely to be pervasive in the published literature.

Here we present an integrated model that accounts for survivorship bias when estimating migratory transitions from geolocators. In a migratory population, the average survival probability of the population is the marginal survival probability across all non-breeding regions. In other words, the survival probability measured by capture-mark-recapture (CMR) methods is the average survival of each non-breeding region weighted by the probability that an individual migrates to each region. As a result, individuals migrating to low-survival regions result in both missing geolocator recoveries and lower survival probability for the population as a whole. Our approach therefore works by integrating the geolocator recovery data with CMR data in a single, unified analysis. An additional benefit of this integrated model is that it also provides estimates of regional non-breeing survival probabilities without the need to collect additional data during the non-breeding season. Our objectives are twofold. First, we use simulated data to demonstrate that the method is able to return unbiased estimates of transition probabilities and to determine how different sampling designs influence estimability of transition probabilities. Second, we apply the model to tracking data from Painted Buntings (*Passerina ciris*), a declining migratory songbird that is thought be threatened by illegal pet trade in parts of its non-breeding range. The estimated transition probabilities and non-breeding survival probabilities from our analysis are consistent with predictions about where Painted Buntings are most at risk of capture during the winter, underscoring the potential of these methods to improve inference from geolocators and reveal new insights into the ecology and conservation of migratory species.

## Materials and methods

We assume that researchers deploy archival geolocators to determine migration routes used by different populations of a focal species with an annual cycle consisting of stationary breeding and non-breeding periods, separated by annual migrations. Researchers deploy geolocators at *s* = 1, 2, 3, …, *S* breeding sites over *t* = 1, 2, 3, …, *T* years and each recovered geolocator is used to assign individuals to one of *j* = 1, 2, …, *J* distinct non-breeding regions. The objective of the study is to determine the proportion of individuals from breeding site *s* that migrate to non-breeding region *j*, which we will represent by a *S* × *J* transition matrix *ψ*. In addition to deployment of geolocators, we assume that researchers also apply marks (e.g., leg bands) to individuals in each breeding site to estimate apparent annual survival at each of the *S* breeding sites using mark-recapture or mark-re-sight methodologies. The integrated model we outline below assumes that geolocator individuals are not included in the mark-capture data set, though small violations of this assumption are unlikely to have practical effects on inference (Abadi, Gimenez, Arlettaz, & Schaub, 2010).

These data provide the following summaries:

- ***N*** : a *S* × *T* matrix containing the number of geolocators deployed at site each breeding site in each year
- ***w***: a *S* × *J* × *T* matrix indicating the number of recovered geolocators from site *s* that spent the non-breeding season in region *j* in year *t*
- ***y**_s_*: annual encounter histories of marked birds at site *s*

For the purposes of this paper, we assume no uncertainty in determining the non-breeding region of each individual, though it may be possible to relax this assumption (see discussion). In most applications, survival data will be collected over longer time scales than geolocator data, which should not pose problems as long as the estimated survival probabilities apply to individuals tracked using geolocators.

Both the geolocator data and the mark-recapture data contain information about the underlying transition matrix *ψ*, allowing us to integrate these two data sets within a single analysis. In the sections below, we describe sub-models for the geolocator recovery and mark-recapture data that allow us to parameterize each model in terms of the underlying transition matrix.

### Geolocator recovery model

Rather than interpret the ***w*** entries as proportional to *ψ* (the default of most archival tagging studies), we derive an explicit geolocator recovery model that treats ***w*** as a random variable governed by both the transition matrix and annual survival probabilities for birds wintering in each non-breeding region. For each breeding site *s*, we model the true number of individuals that went to each non-breeding region *j*:

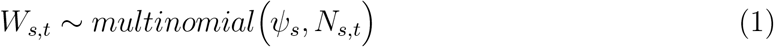

where *W_s,t_* and *s* are vectors of length *J* and 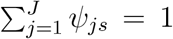. Because non-breeding regions are only observed for individuals that survive and are recaptured, *W_s,t_* is treated as a partially-observed latent parameter. The observed number of geolocator individuals from breeding site *s* that went to non-breeding region *j* can then be modeled as:

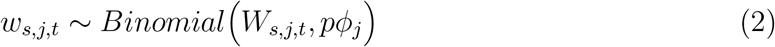

where *ϕ_j_* is the annual apparent survival probability for an individual that spent the non-breeding season in region *j* and *p* is the probability of recapturing a geolocator individual given that it survived and returned. For the model described here, we assume that *p* is constant across all sites and years, though this assumption could be relaxed by including occasion-specific sampling covariates via a logit-link. We also assume *ψ* and *ϕ_j_* are constant across years. This assumption could also be relaxed by including covariates (e.g. sex) and/or allowing temporal variation in one or both parameters. Finally, our calculation of *ϕ_j_* assumes that mortality can occur anywhere during the annual cycle (e.g., on fall migration or on winter quarters) and is independent of breeding location.

### Mark-recapture model

Encounter data from marked individuals can be used to estimate the probability that an individual breeding at site *s* survives and returns to breed the next year (Φ_*s*_)), which is equivalent to the marginal probability of survival across the entire non-breeding area (i.e., the average non-breeding survival weighted by the transition probabilities to each region). Apparent survival can be estimated using a variety of capture-recapture methods, for example the standard Cormack-Jolly-Seber (CJS) model:

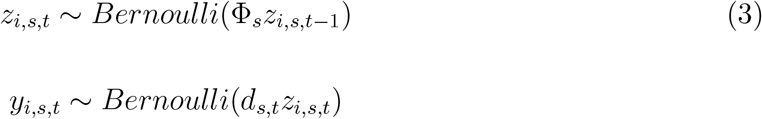

where *z_i,s,t_* is the true state (dead or alive) of individual *i* in year *t* and *d_s,t_* is the probability of detecting an individual given that it is alive and present at site *s* in year *t*. Again, we can model heterogeneity in these parameters by including covariates on detection probability using a logit-link.

Assuming that the marked and geolocator birds have the same survival probability, overall survival probability at each breeding site is equivalent to the average of the non-breeding survivals weighted by the proportion of individuals that spent the non-breeding season in each region:

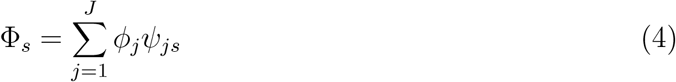

Equation 4 allows us to parameterize the CJS model in terms of the underlying transition matrix *ψ* and thereby integrate the two sub-models.

### Simulations

We used simulated data to determine what biological scenarios yield unbiased estimates of ψ and *ϕ_j_*. In CMR models, parameter identifiability can be assessed by simulating capture histories for a very large number of individuals and then quantifying the bias of parameter estimates from the model (Gimenez, Viallefont, Catchpole, Choquet, & Morgan, 2004). With large sample sizes, the observed frequencies should be equal to their expected values (i.e., no sampling error) and thus any bias in the estimated parameters indicates undentifiabile parameters.

### Identifiability of *ψ* and *ϕ_j_*

To determine whether *ψ* and *ϕ_j_* are identifiable using the model described by equations 1–4, we simulated geolocator recovery and CMR data assuming 10 000 geolocators deployed at each of three breeding sites and 20 000 new individuals added to the CMR data in each year at each site. These values were chosen to be large enough that estimates of *ψ* and *ϕ_j_* were not influenced by sampling error (Gimenez et al., 2004). For each breeding site, we simulated random transition probabilities to each of three non-breeding regions by drawing random values from *Uniform*(0.2, 0.9) and then scaling to ensure the transition probabilities summed to 1. Restricting values to 0.2 − 0.9 ensured that transition probabilities were not close to 0. Transition probabilities for each breeding site were then combined to create the true *ψ* matrix for the simulation. We next simulated a random survival probability for each non-breeding region by sampling from *Uniform*(0.2, 0.9), which spans realistic non-breeding survival probabilities for small migratory birds. Finally, we used equations 1–4 to generate geolocator recoveries and capture histories using the *ψ* matrix and *ϕ* vector created in the previous steps. We repeated these steps 500 times to generate a wide range of transition probabilities and survival probabilities. In all simulations, we assumed 5 years of CMR data, *p* = 0.8 and *d* = 0.6.

For each simulated data set, we estimated the joint likelihood of the model using JAGS version 3.3.0 (Plummer, 2012) called from program R version 3.6.0 (R Core Team, 2016) with package jagsUI version 1.4.2 (Kellner, 2016). Tag recovery probability *p*, detection probability *d*, and non-breeding survival probabilities *ϕ* were given uninformative *Uniform*(0, 1) priors. Priors for the elements of *ψ* were given uninformative Dirichlet priors to ensure that the rows of *ψ* summed to 1. Capture histories were summarized using the multi-dimensional array format for computational efficiency. For all models, we ran three chains for 25,000 iterations each after an adaptation phase of 1,000 iterations and discarding the first 2,000 iterations as burn-in. Convergence was confirmed through Rhat values and visual inspections of trace plots. Code to simulate data and fit the integrated model a re included in Appendix S1.

Denoting the estimated transition probability from site *s* to region *j* for the *i^th^* simulation (*i* ∈ 1 − 500) as 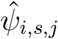, we measured relative bias in the transition estimates to each non-breeding as 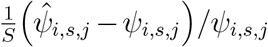. To compare the estimates from the integrated model to the uncorrected estimates from the raw geolocator recoveries, we also converted the elements of ***w***_*i*_ to proportions and measured relative bias in the connectivity to each non-breeding region as 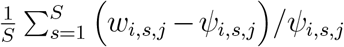. We predicted that the uncorrected estimates would be negatively biased for regions with low survival (transition probabilities underestimated) and positively biased for regions with high survival (transition probabilities overestimated) whereas the corrected estimates from the integrated model would be unbiased for all survival estimates. To test this prediction, we also estimated the correlation between bias of each element of ***w***_*i,s,j*_ and 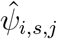 and the corresponding *ϕ_j_* across all simulations. We predicted that the correlation should be positive for the uncorrected estimates and close to 0 for the corrected estimates. Finally, we measured relative bias for each estimate of *ϕ_j_* as 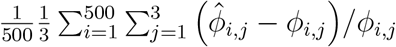

In addition to estimating transition and survival probabilities, many tracking studies are interested in quantifying the strength of migratory connectivity, i.e., the extent to which individuals and populations remain together between seasons of the annual cycle (Cohen et al., 2018). The strength of migratory connectivity, hereafter *MC*, is a function of the transition matrix *ψ*. Estimating *MC* from the uncorrected transition matrix ***w*** could therefore result in biased estimates. However, *MC* measures the relative links between breeding and non-breeding populations and may not suffer from survivorship bias if transition probabilities are equally biased for all focal populations (J. Hostetler *pers comm*). For each simulated data set *i*, we used the `calcMC` function from the `MigConnectivity` R package (Hostetler & Hallworth, 2018) to estimate *MC* using the “true” transition matrix *ψ*(hereafter *MC_ψ,i_*), the raw observation matrix ***w*** (hereafter *MC_w,i_*), and the estimated matrix 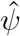 from the integrated model (hereafter 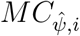). We measured relative bias in *MC_w,i_* and 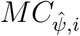 as (*MC_k,i_* − *MC_ψ,i_*)/*MC_ψ,i_*, where *k* equals either *w* or 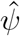, and report the median bias of the 500 simulations.

### Estimability of *ψ*

Even if *ψ* is intrinsically identifiable, a number of factors may influence whether estimates of transition probabilities are unbiased under real-world sampling scenarios, i.e., whether the parameters are estimable (Auger-Méthé et al., 2016). We conducted 3 simulation scenarios to determine what factors influence the estimability of *ψ*:

#### Number of geolocators

To investigate how the number of deployed geolocators and marks influences estimability of *ψ*, we simulated data with sample sizes more typical of geolocator and CMR studies. Specifically, we simulated data assuming 10, 20, 30, 40, and 50 geolocators deployed at each breeding site and 40, 75, 125, 200, 300 individuals added to the CMR data set in each year. As in the identifiability simulations, we assumed 3 breeding and 3 non-breeding sites, 5 years of CMR data, *p* = 0.8 and *d* = 0.6. The number of geolocator and number of marked individuals were varied in a factorial design, resulting in 25 simulation scenarios. For each scenario, we simulated 500 data sets with randomly generated *ψ* matrices and *ϕ_j_* values. *Number of years:* The number of years included in the CMR study may influence estimability of *ψ* by influencing the precision of the ϕ_*s*_ estimates. We simulated data assuming 4, 6, 8, 10, and 12 years of CMR data. As for the other simulations, we generated 500 data sets with random *ψ* matrices and *ϕ_j_* values for each simulation. For all simulations in this scenario, we assumed 3 breeding and 3 non-breeding sites, 30 geolocator deployed at each site, 100 individuals added to the CMR data set in each year, *p* = 0.8 and *d* = 0.6.

#### Number of sites

Increasing or decreasing the number of deployment sites and non-breeding regions could influence estimability through its influence on the precision of survival and transition probability estimates. We simulated data assuming 3, 4, 5, 6, and 7 breeding sites and non-breeding regions. We generated 500 data sets with random *ψ* matrices and *ϕ_j_* values for each simulation and we assumed 5 years of CMR data, 30 geolocators deployed at each site, 100 individuals added to the CMR data set in each year, *p* = 0.8 and *d* = 0.6.

For all estimability simulations, model fitting and evaluation was conducted as described above.

### Application to eastern Painted Bunting population data

Painted Buntings are small (~16g) migratory songbirds that breed in two distinct populations within the United States (Herr, Sykes, & Klicka, 2011). The “western” population breeds primarily in Texas, Oklahoma, Louisiana, and Arkansas while the “eastern” population is restricted to a narrow band of habitat along the coasts of Florida, Georgia, South Carolina, and North Carolina. Monitoring data indicate that the eastern population has declined steadily over the past half century (Sykes Jr. & Holzman, 2005), likely due to habitat loss and the illegal pet trade (Sykes, Holzman, & Iñigo-Elias, 2007; Sykes et al., 2019). Capture of adult males for the pet trade is thought to be a particular problem in Cuba, where keeping wild birds as pets has a long history. However, data on the pet trade are largely anecdotal and the effects of this threat on the demography of Painted Bunting populations have not been demonstrated (Sykes Jr., Manfredi, & Padura, 2006).

In 2017 and 2018, we deployed 180 light-level geolocators (model P50Z11-7-DIP, Migrate Technology Ltd, Coton, Cambridge,UK) on adult male Painted Buntings at 6 sites that span the latitudinal distribution of the eastern population: Carolina Beach State Park, N.C (34.15 N, −77.88 W), Dewee’s Island SC (32.84 N, −79.72 W), Kiawah Island SC (32.61 N, −80.15 W), Spring Island SC (32.33 N, −80.83 W), Little St. Simons GA (31.29 N, −81.34 W), and Little Talbot Island FL (30.46 N, −81.42 W). For the purposes of this analysis, we treated the three South Carolina sites as a single population unit due to their close proximity. Buntings were trapped at feeders using mist nets and traps baited with untreated, white proso millet (*Panicum vergi*). Geolocators were attached with a leg-loop harness (Rappole and Tipton 1991). We recovered 65 geolocators in 2018 and 2019; 61 had viable data. Recovery effort was similar among all sites. We used the R package `SGAT` (Lisovski & Hahn, 2012) to generate location estimates from the raw light data. Twilights were identified using the function `preprocessLight` from the R package `TwGeos` (Lisovski, Wotherspoon, & Sumner, 2016). The appropriate user-defined threshold level varied by individual and ranged from 1-11.5. We used `SGAT` to determine appropriate zenith angles for each bird during the stationary breeding period when individuals are at known locations. To determine appropriate zenith angles at times of the year when location is unknown (i.e., the non-breeding season), we used the Hill-Ekstrom calibration method implemented using the function `findHEZenith` from the R package `TwGeos` (Lisovski et al., 2016). For each individual, we took a weighted median of all generated locations during the core of the wintering period (December and January) and used that centroid to define each individual’s wintering location. Individuals were then assigned to one of the following four regions based on their estimated wintering centroid: northern Florida, southern Florida, the Bahamas, or Cuba (Table S1).

Survival data for this analysis were collected from 1999$-$2005 as part of a separate study on the demography of eastern Painted Buntings (Sykes et al., 2019). Buntings were captured in mist nests situated at feeders filled with millet. Captured birds were immediately removed from mist nets, aged (hatch year, second-year, or after-second-year), sexed, and banded. Birds were marked with three colored plastic leg bands and one US Geological Survey (USGS) numbered aluminum leg band arranged 2 on each leg, in unique 4-band combinations for individual identification. Unbanded individuals were marked through 2003. In subsequent years, missing or faded color-bands were replaced on individuals still retaining the USGS metal bands, when possible. Banded buntings were re-sighted at feeders during observation sessions starting in 2001. Observation sessions were conducted once annually during the breeding season at each site. In years when birds were banded (2001 − 2003), observation sessions were conducted at least one day before banding sessions. Feeders were arranged so all open feeding ports were visible by observers using 20-60x zoom spotting scopes, approximately 10m away from the feeders. See Sykes et al. (2019) for further details. For this analysis, we included only data from males originally banded as adults (either second-year or after-second-year; n = 402). We further included only banding locations within 20km of our geolocator deployment sites to ensure that survival estimates from the re-sight data corresponded as closely as possible to the geolocator study (Table S2).

Geolocator recovery and CMR data were used to parameterize the integrated model. Mark re-sight data from individual feeders (Table S2) were pooled to estimate a single survival probability for each region (North Carolina, South Carolina, Georgia, and Florida), forming the basis for implementing Eq. 4 in this analysis. Thus, we did not model annual variation in survival and assumed that the time-averaged survival estimates from the mark-re-sight data are representative of the survival probabilities experienced by birds in the geolocator study. Following Sykes et al. (2019), we included site-specific effects of human development within 700m and feeder-level random effects on survival. We also included feeder-specific effort covariates in the detection model (Sykes et al., 2019). Using apparent survival estimates, we estimated weighted transition probabilities from each breeding region to each wintering region. To examine differences in biased and unbiased transition probabilities, we calculated unweighted (raw) transition probabilities by dividing the number of geolocators recovered at a breeding site from a single wintering region by the total number of geolocators recovered at a site from any wintering region. Code and data needed to implement the integrated model are included in Appendix S1. We used uninformative priors for all parameters. Posterior distributions were based on 3 chains, run for 20000 iterations each after discarding the first 2500 iterations as burn-in. Model convergence was assessed using R-hat values (Brooks & Gelman, 1998) and by visual inspection of trace plots.

## Results

### Simulation

Both *ψ* and *ϕ_j_* were identifiable using the integrated model (Fig. 1). As predicted, the correlation between non-breeding survival and the bias in the uncorrected estimates of ψ was positive (1.27), signifying that the transition probabilities for low survival regions were underestimated and overestimated for high survival regions. This structural bias was substantial for the extreme low and high survival regions (*ϕ_j_* = 0.2 or 0.9), under- and overestimating *ψ*_j_ by nearly 45%, respectively. In contrast, the integrated model returned unbiased estimates of *ψ* regardless of the underlying survival probabilities (correlation = −0.058), leading to minimal relative bias < 1% even at the most extreme low and high survival sites. Estimates of *ϕ_j_* from the integrated model were also minimally biased (0.017). Estimates of *MC* showed no bias when using the transition matrix from the integrated model (median relative bias < 0.0001) and a small negative bias for the uncorrected estimates (median relative bias = −0.04). Although bias in *MC_w_* was small on average, these estimates showed much larger variation than the 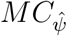 estimates (Fig. 2). Root-mean-square error of the *MC_w_* estimates was 0.03, nearly 10 times higher than the root-mean-square error of the 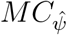 estimates (0.004).

**Figure 1:**
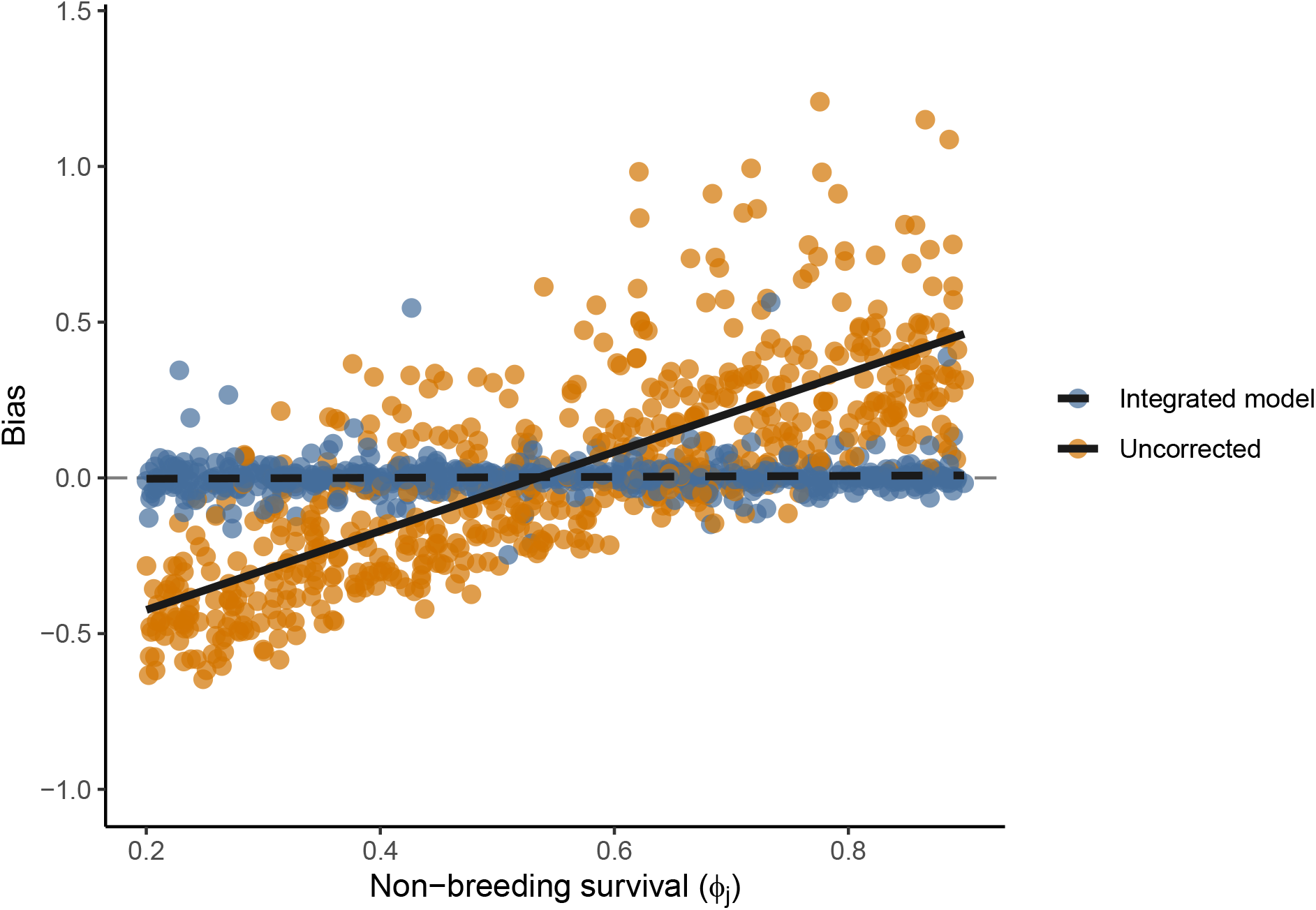
Correlation between non-breeding survival and bias in the estimated transition probabilities for identifiability simulations

**Figure 2:**
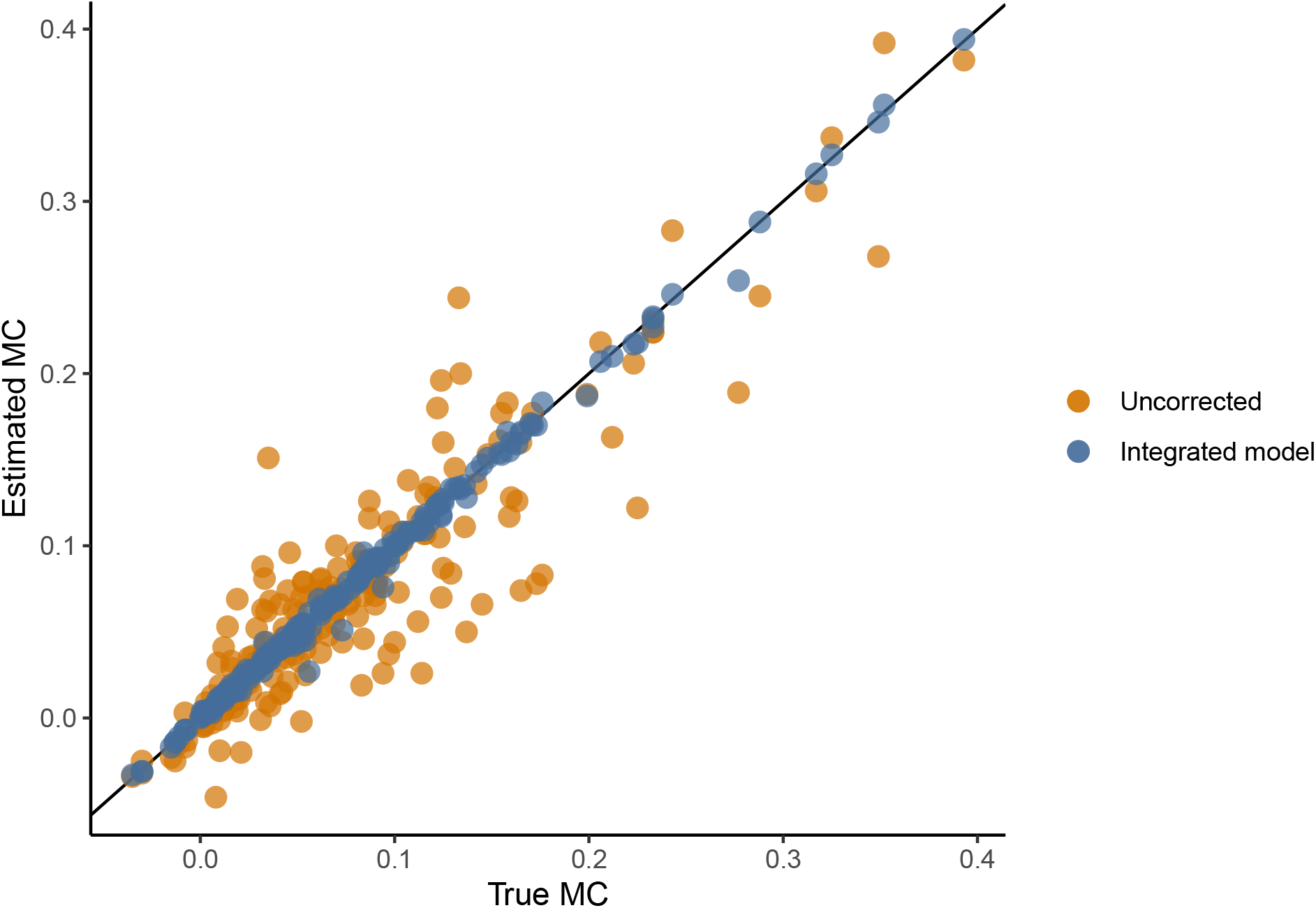
Relationship between the true and estimated strength of migratory connectivity for the 500 identifiability simulations. Estimated MC are based on either the uncorrected transition probabilities (orange points) or the transition probabilities from the integrated model (blue points). The black line indicates a 1:1 relationship between the true and estimated MC values.

Estimability simulations indicated that, under more realistic sample sizes, some bias remains in the estimates of *ψ* and *ϕ_j_*, though the integrated model greatly reduced bias in ψ relative to the uncorrected estimates. Across all scenarios, relative bias of *ψ_j_* for the extreme low and high survival regions was generally < 10% for the integrated model, much lower than the ∼ 40 − 50% of the uncorrected estimates. Even with only 10 geolocators deployed at each site and 75 individuals per year added to the capture-recapture data (Fig. 3), the correlation between bias and non-breeding survival was much lower in the integrated model (0.51, 0.46 − 0.56) relative to the uncorrected estimates (1.27, 1.2 − 1.33). Deploying additional geolocators reduced bias substantially, whereas adding additional individuals to the CMR data set had less influence on bias (Table 1). Bias in the integrated model generally decreased as both the length of the CMR study and the number of breeding sites and non-breeding regions increased (See Appendix S2), though the decreases in bias were small and the value of adding more years/sites generally decreased as more were added. Estimates of *ϕ_j_* were positively biased by ∼ 2 − 5% under all estimability scenarios. Bias in *ϕ_j_* generally decreased as more marks (Table 1) and more years of CMR data were included in the analysis but was not sensitive to the number of sites (Appendix S2).

**Table 1:**
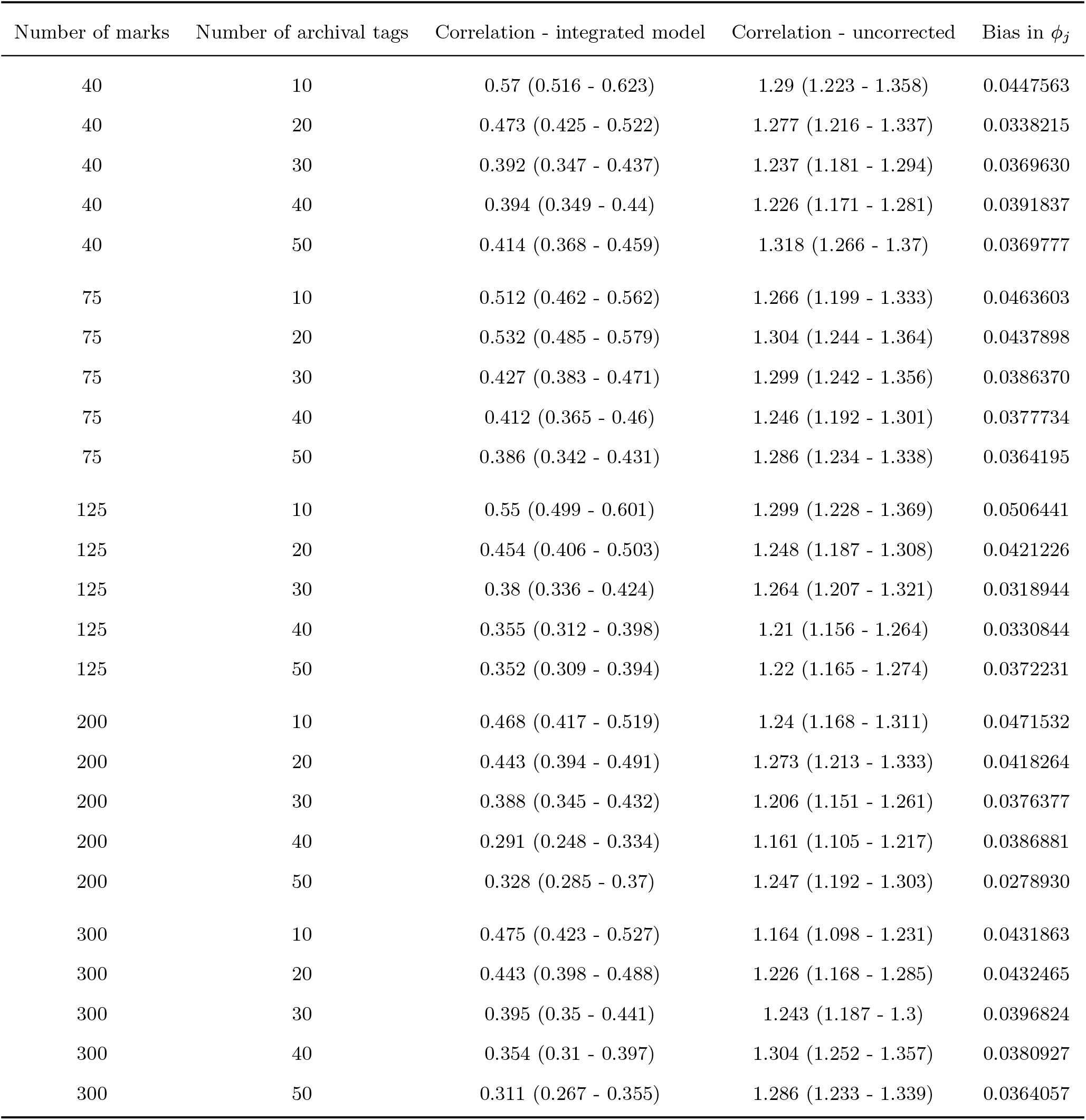
Estimability of transition probabilities and non-breeding survival probabilities as a function of the number of marks and geolocators deployed. Correlations are the mean and 95% confidence interval from the 500 simulated data sets

| Number of marks | Number of archival tags | Correlation - integrated model | Correlation - uncorrected | Bias in $\phi_j$ |
| --- | --- | --- | --- | --- |
| 40 | 10 | 0.57 (0.516 - 0.623) | 1.29 (1.223 - 1.358) | 0.0447563 |
| 40 | 20 | 0.473 (0.425 - 0.522) | 1.277 (1.216 - 1.337) | 0.0338215 |
| 40 | 30 | 0.392 (0.347 - 0.437) | 1.237 (1.181 - 1.294) | 0.0369630 |
| 40 | 40 | 0.394 (0.349 - 0.44) | 1.226 (1.171 - 1.281) | 0.0391837 |
| 40 | 50 | 0.414 (0.368 - 0.459) | 1.318 (1.266 - 1.37) | 0.0369777 |
| 75 | 10 | 0.512 (0.462 - 0.562) | 1.266 (1.199 - 1.333) | 0.0463603 |
| 75 | 20 | 0.532 (0.485 - 0.579) | 1.304 (1.244 - 1.364) | 0.0437898 |
| 75 | 30 | 0.427 (0.383 - 0.471) | 1.299 (1.242 - 1.356) | 0.0386370 |
| 75 | 40 | 0.412 (0.365 - 0.46) | 1.246 (1.192 - 1.301) | 0.0377734 |
| 75 | 50 | 0.386 (0.342 - 0.431) | 1.286 (1.234 - 1.338) | 0.0364195 |
| 125 | 10 | 0.55 (0.499 - 0.601) | 1.299 (1.228 - 1.369) | 0.0506441 |
| 125 | 20 | 0.454 (0.406 - 0.503) | 1.248 (1.187 - 1.308) | 0.0421226 |
| 125 | 30 | 0.38 (0.336 - 0.424) | 1.264 (1.207 - 1.321) | 0.0318944 |
| 125 | 40 | 0.355 (0.312 - 0.398) | 1.21 (1.156 - 1.264) | 0.0330844 |
| 125 | 50 | 0.352 (0.309 - 0.394) | 1.22 (1.165 - 1.274) | 0.0372231 |
| 200 | 10 | 0.468 (0.417 - 0.519) | 1.24 (1.168 - 1.311) | 0.0471532 |
| 200 | 20 | 0.443 (0.394 - 0.491) | 1.273 (1.213 - 1.333) | 0.0418264 |
| 200 | 30 | 0.388 (0.345 - 0.432) | 1.206 (1.151 - 1.261) | 0.0376377 |
| 200 | 40 | 0.291 (0.248 - 0.334) | 1.161 (1.105 - 1.217) | 0.0386881 |
| 200 | 50 | 0.328 (0.285 - 0.37) | 1.247 (1.192 - 1.303) | 0.0278930 |
| 300 | 10 | 0.475 (0.423 - 0.527) | 1.164 (1.098 - 1.231) | 0.0431863 |
| 300 | 20 | 0.443 (0.398 - 0.488) | 1.226 (1.168 - 1.285) | 0.0432465 |
| 300 | 30 | 0.395 (0.35 - 0.441) | 1.243 (1.187 - 1.3) | 0.0396824 |
| 300 | 40 | 0.354 (0.31 - 0.397) | 1.304 (1.252 - 1.357) | 0.0380927 |
| 300 | 50 | 0.311 (0.267 - 0.355) | 1.286 (1.233 - 1.339) | 0.0364057 |

**Figure 3:**
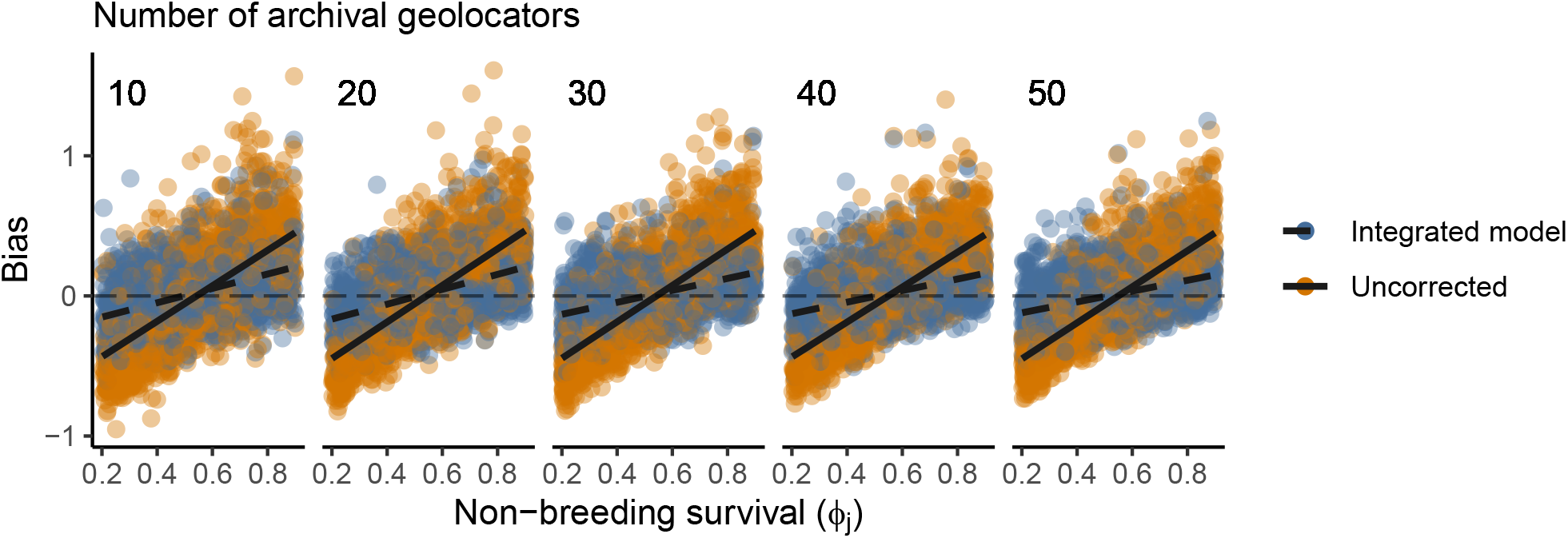
Correlation between non-breeding survival and bias in the estimated transition probabilities as a function of the number of marks and archival geolocators

### Painted Bunting survival and migratory connectivity

Estimates for apparent annual survival within breeding regions were similar for Florida (mean: 0.68, credible interval: 0.58 − 0.76), Georgia (0.68, 0.59 − 0.76), and South Carolina (0.66, 0.58 − 0.74), but were notably lower in North Carolina (0.6, 0.48 − 0.71). Of the 180 geolocators deployed in 2017 and 2018, we recovered 61 geolocators with usable data (Table S1). Raw transition probabilities estimated from these geolocators suggested that Painted Buntings breeding in Florida were relatively evenly distributed among the four non-breeding regions, birds from Georgia and South Carolina were most likely to winter in south Florida, and birds from North Carolina were most likely to winter in Cuba (Table 2). No geolocators were recovered in North Carolina from birds that had wintered in north or south Florida, and therefore transition probabilities to these wintering regions were estimated to be zero based on the raw recoveries.

**Table 2:**
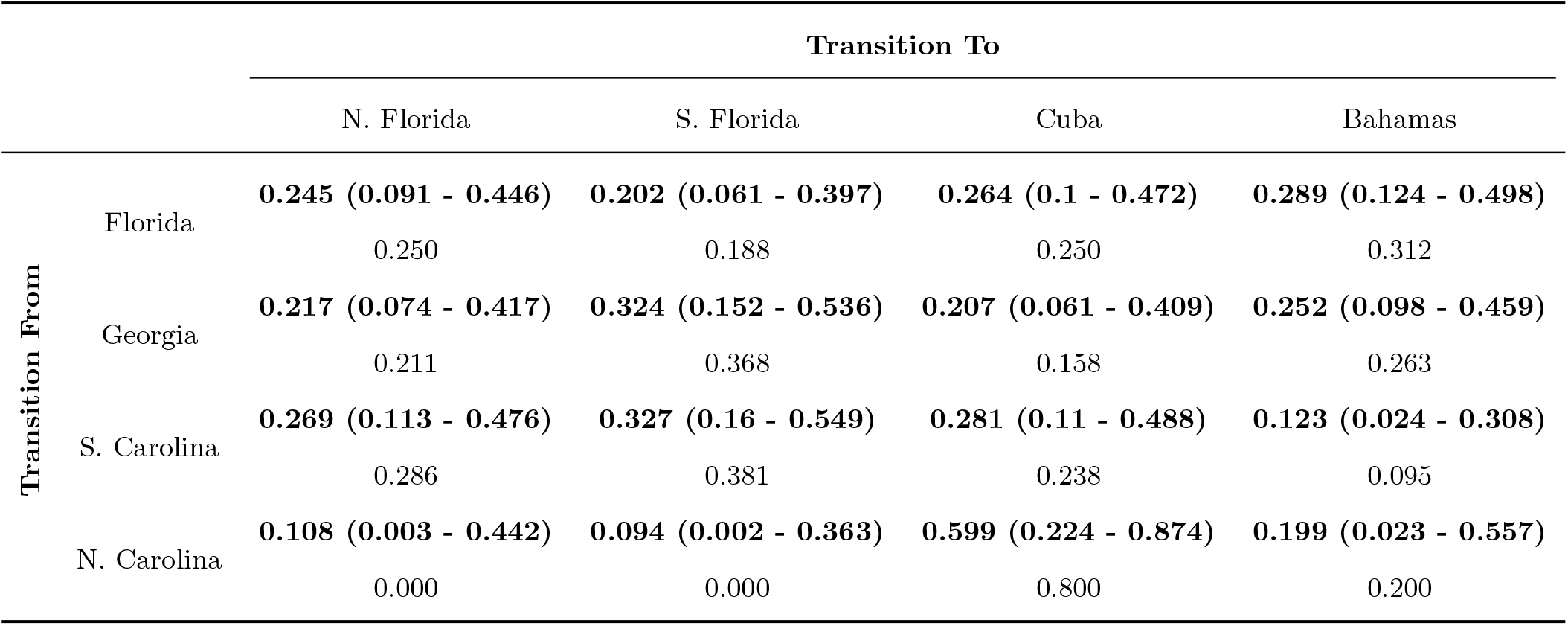
Transition probabilities from breeding sites to wintering sites, estimated from geolocators recovered on the breeding grounds in the 2017 and 2018 field seasons. Bolded values are the posterior mean of *ψ* from the integrated models, with estimated 95% credible intervals in parentheses. Non-bolded values are the equivalent transition probabilities estimated from the raw recovery data. Note that the raw *ψ* estimates are point estimates and do not provide any measure of uncertainty.

Results from the integrated model suggest that winter region affected annual survival. Survival was comparable for buntings wintering in north Florida (0.72, 0.4 − 0.98), south Florida (0.75, 0.49 − 0.98), and the Bahamas (0.72, 0.39 − 0.98; Figure 4). Male buntings wintering in Cuba, however, had measurably lower annual survival probabilities (0.57, 0.37 − 0.9). Due to regional survival variation in non-breeding survival, transition probabilities from the integrated model differed from the raw estimates in important ways (Table 2). In particular, the integrated model suggested that birds breeding in Georgia and South Carolina breeding populations were less likely to winter in south Florida and more likely to winter in Cuba than suggested by the raw recovery data. These differences are consistent with over-estimation of connectivity to south Florida due to the high survival in that region and under-estimation of connectivity to Cuba due to the low survival there. For the North Carolina population, the integrated model actually indicated lower connectivity to Cuba than the raw data. This unintuitive result occurred because the integrated model estimated low but non-zero connectivity to north and south Florida despite no geolocator recoveries from these regions. In total, estimated connectivity between North Carolina and Florida was ~20%, which more than offset differences between the integrated and raw connectivity estimates to Cuba.

**Figure 4:**
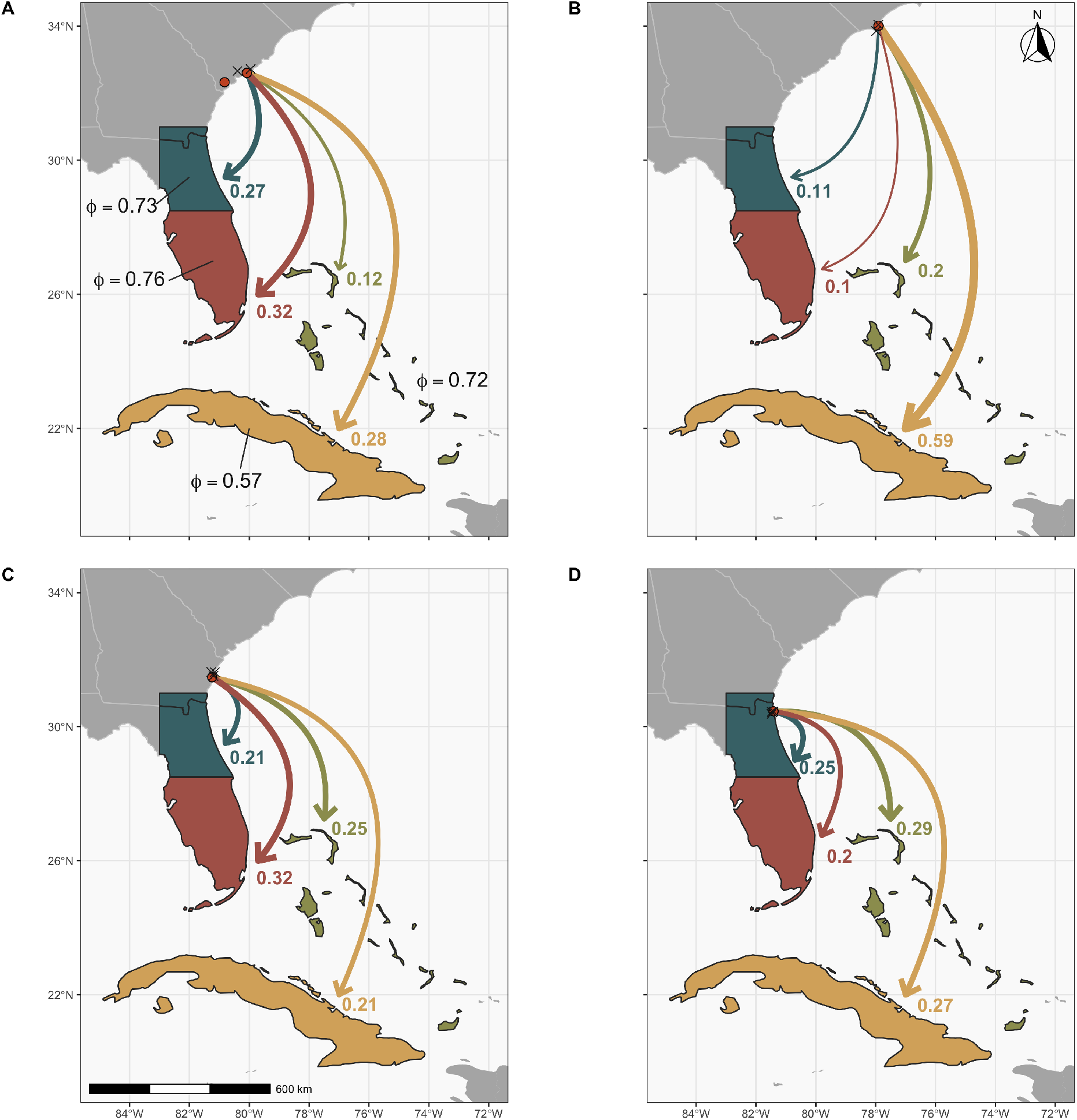
Estimated transition probabilities and non-breeding survival of southeastern Painted Buntings (*Passerina ciris*) from (A) Kiawah Island and Spring Island, SC, (B) Wilmington, NC, (C) Little St. Simons Island, GA, and (D) Big Talbot Island State Park, FL. Red dots indicate geolocator deployment sites and black crosses indicate sites where mark-re-sight data was collected. Estimated survival probabilities for each of the four non-breeding regions are shown in panel A. Values at the base of each arrow are the estimated transition probabilities from the integrated model for each breeding site and non-breeding region. Arrows are color-coded by non-breeding region and sized relative to the estimated transition probabilities.

## Discussion

Archival geolocators have transformed the study of small, migratory organisms but yield biased observations because geolocators can only be recovered from surviving individuals. Our simulations confirm that estimating migratory transition probabilities from observed geolocator recoveries produces biased estimates when survival probabilities differ among non-breeding regions. Given that regional variation in non-breeding survival is the rule rather than the exception for most migratory species (Hewson, Thorup, Pearce-Higgins, & Atkinson, 2016; Healy et al., 2017; Kramer et al., 2018), survivorship bias is likely to be a pervasive issue for studies using archival tracking devices. As we demonstrate, the integration of a probabilistic geolocator recovery model with mark-recapture data can reduce or eliminate survivorship bias, providing a practical approach for strengthening inferences about migratory connectivity.

The use of archival geolocators has grown dramatically over the past decade, with over 120 publications in 2016 and 2017 alone (McKinnon & Love, 2018), motivating multiple research teams to investigate the effects of these geolocators on the survival of tracked individuals (Costantini & Møller, 2013; Peterson et al., 2015; Brlík et al., 2020). However, we are not aware of any study that has been designed to account for survivorship bias on interpretation of data from archival geolocators. Our results indicate that assuming observed geolocator recoveries accurately reflect the underlying migration patterns is potentially problematic and in extreme cases could lead to false inferences about which non-breeding regions are most important for specific breeding populations. Reliance on raw recovery data may be especially problematic for archival tracking studies based on small sample sizes of deployed geolocators. Mortality or dispersal could result in no individuals being observed to migrate to certain non-breeding regions. This was the case for our sample of Painted Buntings from North Carolina, which included no birds wintering in north or south Florida. In this case, the true transition probabilities are unlikely to be 0; instead, these zeros are probably the result of sampling variation with only a small number of tracked birds. Defining an explicit geolocator recovery model that treats the observed data as realizations of one or more stochastic process or sampling models provides a means to account for sampling variation and estimate the underlying process model parameters, even without integrating the survival data in the model.

Integrating geolocator recovery and survival data enables novel insights that neither source of information alone can provide. Our integrated model provided compelling evidence that survival probabilities of Painted Buntings wintering in Cuba may be 20% lower than for birds that winter in other parts of the range. Although the non-breeding survival rates had high uncertainty, these results are consistent with the hypothesis that Painted Buntings wintering in Cuba are at higher risk of being captured as part of the illegal pet trade than birds wintering elsewhere. Reliable estimates of the number of buntings captured in Cuba or Florida are not available but anecdotal reports have documented localized trapping of as many as 700 buntings over three days in Cuba (Sykes Jr. et al., 2006; Sykes et al., 2007). Additionally, of the 80 geolocators deployed in 2018 as part of this study, at least two are known to have been captured in Cuba and Sykes et al. (2007) confirmed 19 buntings banded in the US and captured in Cuba. Of course, other explanations could explain our results, including longer migration distances required to reach Cuba or differences in habitat loss throughout the winter range. Our results nonetheless demonstrate how integrating tracking and survival data can lead to deeper insights about the dynamics of migratory species.

A common objective of recent tracking studies is to estimate the strength of migratory connectivity (Cohen et al., 2018), an index of the extent that individuals and populations remain together between seasons of the annual cycle. Even when the underlying transition probabilities are biased due to differential survival, our simulations indicate that *MC* is largely unbiased when derived from observed geolocator recoveries. This result is not unexpected because the *MC* metric is based on the relative links between breeding and non-breeding populations, rather than the absolute transition probabilities. In other words, transition probabilities to low-survival, non-breeding regions will be underestimated for all breeding sites and as a result the correlation of distances between breeding and non-breeding regions will remain unbiased. These results suggest that, on average, published estimates of *MC* based on archival geolocators should be considered unbiased. However, we found that *MC* estimates based on the integrated model were, on average, more precise, i.e. much closer to the true *MC* value, than estimates based on raw recovery data. This result is likely due to explicitly accounting for process and sampling uncertainty in the geolocator recovery model. By viewing the realized recoveries as stochastic processes, the integrated model attempts to separate the expected transition and survival probabilities from the process and sampling noise inherent to the data collection process. Reducing the influence of sampling and process uncertainty in the *ψ* estimates in turn reduces noise in the estimates of *MC* compared to estimates based on the raw geolocator recoveries. Future work that allows researchers to directly account for survivorship bias and produce more accurate estimates of *MC* would be beneficial.

Our integrated approach permits unbiased estimates of non-breeding survival and transition probabilities for seasonal ranges, but makes several assumptions. First, we assume that non-breeding survival is independent of breeding origin. If non-breeding survival differs among breeding sites, because for example individuals migrate different distances, then site-specific transition probabilities will be confounded with variation in site-specific non-breeding survival. Second, individuals tracked using archival geolocators are assumed to have the same survival probability as individuals in the capture-recapture data set. Recent meta-analyses of light-level geolocator studies found that these geolocators can have a small, negative effect on survival, especially for small and highly aerial species (Costantini & Møller, 2013; Brlík et al., 2020). However, geolocator effects should not bias transition probability estimates as long as they are independent of where individuals overwinter. In this case, the lower return rate of geolocator individuals will be reflected in lower geolocator recovery probabilities but other parameters should be unbiased. Third, integrating geolocator recovery and capture-recapture models assumes independence of the two data sets. Strictly speaking, if some individuals are shared between the two data sets, variance parameters in the model will be underestimated, though small violations of this assumption are unlikely to affect inference (Abadi et al., 2010). Fourth, we assumed no uncertainty in the non-breeding region of each tracked individual. Some archival geolocators, particularly light-level geolocators, have substantial uncertainty and even with large non-breeding regions, some individuals may not be assigned to regions with 100% accuracy. Uncertainty in non-breeding regions could be incorporated into the analysis by treating non-breeding assignments as categorical random variables with probabilities estimated from the raw tracking data, though this modification is beyond the scope of this paper. Fifth, assumptions of the chosen survival model also apply to the integrated geolocator recovery model.

Prior to the development of miniaturized archival tracking devices, migratory animals could not be tracked across their entire annual cycle. This technology has created novel research opportunities on thousands of species, but data from these geolocators must be carefully interpreted. Discussions about inferences from archival geolocators has mainly focused on whether geolocators influence the fitness or behavior of tracked individuals (Arlt, Low, & Pärt, 2013; Costantini & Møller, 2013; Brlík et al., 2020). Although these issues are important, the effects of survivorship bias on inference from archival geolocators have not been widely acknowledged. Our results demonstrate that survivorship bias, a potentially ubiquitous outcome of archival geolocators, can be reduced or eliminated if data for estimating survival is also available for each deployment site. Thanks to the focus on quantifying geolocator effects, many tracking studies are likely already collecting mark-recapture or mark-re-sight data that could be used to fit the integrated model presented here. In cases where researchers are designing new tracking studies, collection of these auxiliary survival data should be prioritized both to measure geolocator effects and obtain accurate transition probabilities and regional non-breeding survival probabilities. Improving inferences from archival geolocators through proper data collection and further development of integrated models will enable this technology to further transform the study of small, migratory organisms.

## Supporting information

Code for simulations, case study, and figures

## Acknowledgments

We thank the many volunteers who maintained feeders and reported Painted Bunting encounters, and the Spring Island Trust, Bald Head Island Conservancy, Little St. Simons Island, and North Florida Land Trust for providing housing and support during geolocator deployment and recovery. Thanks to J.A. Royle and J. Sauer for comments that improved earlier drafts of this paper. This work was supported by a Neotropical Migratory Bird Protection Act grant (#6759) from the U.S. Fish & Wildlife Service and by funding from the Disney Conservation Fund, the Spring Island Land Trust, the Cornell Lab of Ornithology, the Smithsonian Institution, and Utah State University. Use of trade, product, or firm names does not imply endorsement by the U.S. Government.

## Author contributions

CSR conceived the study and conducted simulations. CSR, AJS, VRG, AMG, and TSS designed and conducted the geolocator study. PWS and MCF designed and carried out the mark-re-sight study. AJS analyzed the geolocator data and AVT produced the integrated analysis. CSR, VRG, MCF, PWS, and TSS acquired funding. CSR, AJS, and AVT wrote the first version of the manuscript and all authors contributed to subsequent revisions.

## Animal welfare

All research activities were performed under protocols approved by the animal care and use committees of the authors’ institutions, permitted by the relevant federal and state authorities, and in compliance with the Guidelines to the Use of Wild Birds in Research (Fair, Paul, and Jones, 2010).

## Data availability

All data associated with this study will be deposited in a Dryad repository upon acceptance. Code to carry out the simulations and the case study analysis can be found in Appendix S1.

## References

Abadi, F., Gimenez, O., Arlettaz, R., & Schaub, M. (2010). An assessment of integrated population models: Bias, accuracy, and violation of the assumption of independence. Ecology, 91 (1), 7–14. doi:10.1890/08-2235.1

Arlt, D., Low, M., & Pärt, T. (2013). Effect of Geolocators on Migration and Subsequent Breeding Performance of a Long-Distance Passerine Migrant. PLoS ONE, 8 (12). doi:10.1371/journal.pone.0082316

Auger-Méthé, M., Field, C., Albertsen, C. M., Derocher, A. E., Lewis, M. A., Jonsen, I. D., & Flemming, J. M. (2016). State-space models’ dirty little secrets: Even simple linear Gaussian models can have estimation problems. Scientific Reports, 6 (1), 1–10. doi:10.1038/srep26677

Brlík, V., Koleček, J., Burgess, M., Hahn, S., Humple, D., Krist, M., … Procházka, P. (2020). Weak effects of geolocators on small birds: A meta-analysis controlled for phylogeny and publication bias. Journal of Animal Ecology, 89 (1), 207–220. doi:10.1111/1365-2656.12962

Brooks, S. P., & Gelman, A. (1998). General Methods for Monitoring Convergence of Iterative Simulations. Journal of Computational and Graphical Statistics, 7 (4), 434–455. doi:10.1080/10618600.1998.10474787

Brownie, C., Hines, J. E., Nichols, J. D., Pollock, K. H., & Hestbeck, J. B. (1993). Capture-Recapture Studies for Multiple Strata Including Non-Markovian Transitions. Biometrics, 49 (4), 1173–1187. doi:10.2307/2532259

Cohen, E. B., Hostetler, J. A., Hallworth, M. T., Rushing, C. S., Sillett, T. S., & Marra, P. P. (2018). Quantifying the strength of migratory connectivity. Methods in Ecology and Evolution, 9 (3), 513–524. doi:10.1111/2041-210X.12916

Cohen, E. B., Hostetler, J. A., Royle, J. A., & Marra, P. P. (2014). Estimating migratory connectivity of birds when re-encounter probabilities are heterogeneous. Ecology and Evolution, 4 (9), 1659–1670. doi:10.1002/ece3.1059

Cooper, N. W., Ewert, D. N., Jr, J. M. W., Helmer, E. H., & Marra, P. P. (2019). Revising the wintering distribution and habitat use of the Kirtland’s warbler using play-back surveys, citizen scientists, and geolocators. Endangered Species Research, 38, 79–89. doi:10.3354/esr00937

Costantini, D., & Møller, A. P. (2013). A meta-analysis of the effects of geolocator application on birds. Current Zoology, 59 (6), 697–706. doi:10.1093/czoolo/59.6.697

Delmore, K. E., Fox, J. W., & Irwin, D. E. (2012). Dramatic intraspecific differences in migratory routes, stopover sites and wintering areas, revealed using light-level geolo-cators. Proceedings of the Royal Society B: Biological Sciences, 279 (1747), 4582–4589. doi:10.1098/rspb.2012.1229

Domeier, M. L., & Nasby-Lucas, N. (2013). Two-year migration of adult female white sharks (Carcharodon carcharias) reveals widely separated nursery areas and conservation concerns. Animal Biotelemetry, 1 (1), 2. doi:10.1186/2050-3385-1-2

Eckert, S. A., & Stewart, B. S. (2001). Telemetry and satellite tracking of whale sharks, Rhincodon typus, in the Sea of Cortez, Mexico, and the north Pacific Ocean. In E. K. Balon, T. C. Tricas, & S. H. Gruber (Eds.), The behavior and sensory biology of elasmobranch fishes: An anthology in memory of Donald Richard Nelson (Vol. 20, pp. 299–308). Dordrecht: Springer Netherlands. doi:10.1007/978-94-017-3245-1_17

Fraser, K. C., Stutchbury, B. J. M., Silverio, C., Kramer, P. M., Barrow, J., Newstead, D., … Tautin, J. (2012). Continent-wide tracking to determine migratory connectivity and tropical habitat associations of a declining aerial insectivore. Proceedings of the Royal Society B: Biological Sciences, 279 (1749), 4901–4906. doi:10.1098/rspb.2012.2207

Gimenez, O., Viallefont, A., Catchpole, E. A., Choquet, R., & Morgan, B. J. (2004). Methods for investigating parameter redundancy. Animal Biodiversity and Conservation, 27 (1), 561–572.

Hallworth, M. T., & Marra, P. P. (2015). Miniaturized GPS Tags Identify Non-breeding Territories of a Small Breeding Migratory Songbird. Scientific Reports, 5 (1), 11069. doi:10.1038/srep11069

Healy, S. J., Hinch, S. G., Porter, A. D., Rechisky, E. L., Welch, D. W., Eliason, E. J., … Furey, N. B. (2017). Route-specific movements and survival during early marine migration of hatchery steelhead Oncorhynchus mykiss smolts in coastal British Columbia. Marine Ecology Progress Series, 577, 131–147. doi:10.3354/meps12238

Herr, C. A., Sykes, P. W., & Klicka, J. (2011). Phylogeography of a vanishing North American songbird: The Painted Bunting (Passerina ciris). Conservation Genetics, 12 (6), 1395–1410. doi:10.1007/s10592-011-0237-6

Hewson, C. M., Thorup, K., Pearce-Higgins, J. W., & Atkinson, P. W. (2016). Population decline is linked to migration route in the Common Cuckoo. Nature Communications, 7 (1), 1–8. doi:10.1038/ncomms12296

Hostetler, J. A., & Hallworth, M. T. (2018). MigConnectivity: Estimate Strength of Migratory Connectivity for Migratory Animals. R package version 0.3.0.

Jiménez López, M. E., Palacios, D. M., Jaramillo Legorreta, A., Urbán R., J., & Mate, B. R. (2019). Fin whale movements in the Gulf of California, Mexico, from satellite telemetry. PLOS ONE, 14 (1), e0209324. doi:10.1371/journal.pone.0209324

Kellner, K. (2016). JagsUI: A wrapper around rjags to streamline JAGS analyses. R package v. 1.4.2.

Korner-Nievergelt, F., Sauter, A., Atkinson, P. W., Guélat, J., Kania, W., Kéry, M., … Noordwijk, A. J. V. (2010). Improving the analysis of movement data from marked individuals through explicit estimation of observer heterogeneity. Journal of Avian Biology, 41 (1), 8–17. doi:10.1111/j.1600-048X.2009.04907.x

Kramer, G. R., Andersen, D. E., Buehler, D. A., Wood, P. B., Peterson, S. M., Lehman, J. A., … others. (2018). Population trends in Vermivora warblers are linked to strong migratory connectivity. Proceedings of the National Academy of Sciences, 115 (14), E3192–E3200.

Lisovski, S., & Hahn, S. (2012). GeoLight – processing and analysing light-based geolocator data in R. Methods in Ecology and Evolution, 3 (6), 1055–1059. doi:10.1111/j.2041-210X.2012.00248.x

Lisovski, S., Wotherspoon, S., & Sumner, M. (2016). TwGeos: Basic data processing for light-level geolocation archival tags. R package v 0.1.2.

McKinnon, E. A., & Love, O. P. (2018). Ten years tracking the migrations of small landbirds: Lessons learned in the golden age of bio-loggingDiez años siguiendo las migraciones de aves terrestres pequeñas: Lecciones aprendidas en la edad de oro de los bio-registrosTracking migration in the golden age of bio-logging. The Auk, 135 (4), 834–856. doi:10.1642/AUK-17-202.1

Naidoo, R., Chase, M. J., Beytell, P., Du Preez, P., Landen, K., Stuart-Hill, G., & Taylor, R. (2016). A newly discovered wildlife migration in Namibia and Botswana is the longest in Africa. Oryx, 50 (1), 138–146. doi:10.1017/S0030605314000222

Nichols, J. D., Reynolds, R. E., Blohm, R. J., Trost, R. E., Hines, J. E., & Bladen, J. P. (1995). Geographic Variation in Band Reporting Rates for Mallards Based on Reward Banding. The Journal of Wildlife Management, 59 (4), 697–708. doi:10.2307/3801946

Peterson, S. M., Streby, H. M., Kramer, G. R., Lehman, J. A., Buehler, D. A., & Andersen, D. E. (2015). Geolocators on Golden-winged Warblers do not affect migratory ecologyLos geo-localizadores no afectan la ecología migratoria de Vermivora chrysopteraGeolocators on small songbirds. The Condor, 117 (2), 256–261. doi:10.1650/CONDOR-14-200.1

Plummer, M. (2012). JAGS: Just Another Gibbs Sampler version 3.3.0.

R Core Team. (2016). R: A Language and Environment for Statistical Computing. Vienna, Austria: R Foundation for Statistical Computing. Retrieved from https://www.R-project.org/

Richter, H. V., & Cumming, G. S. (2008). First application of satellite telemetry to track African straw-coloured fruit bat migration. Journal of Zoology, 275 (2), 172–176. doi:10.1111/j.1469-7998.2008.00425.x

Sawyer, H., Kauffman, M. J., Nielson, R. M., & Horne, J. S. (2009). Identifying and prioritizing ungulate migration routes for landscape-level conservation. Ecological Applications, 19 (8), 2016–2025. doi:10.1890/08-2034.1

Scarpignato, A. L., Harrison, A.-L., Newstead, D. J., Niles, L. J., Porter, R. R., Tillaart, M. van den, & Marra, P. P. (2016). Field-testing a new miniaturized GPS-Argos satellite transmitter (3.5 g) on migratory shorebirds. Wader Study, 123 (3). doi:10.18194/ws.00046

Smith, M., Bolton, M., Okill, D. J., Summers, R. W., Ellis, P., Liechti, F., & Wilson, J. D. (2014). Geolocator tagging reveals Pacific migration of Red-necked Phalarope Phalaropus lobatus breeding in Scotland. Ibis, 156 (4), 870–873. doi:10.1111/ibi.12196

Sykes, P. W., Freeman, M. C., Sykes, J. J., Seginak, J. T., Oleyar, M. D., & Egan, J. P. (2019). Annual survival, site fidelity, and longevity in the eastern coastal population of the Painted Bunting (Passerina ciris). The Wilson Journal of Ornithology, 131 (1), 96–110. doi:10.1676/18-56

Sykes, P. W., Holzman, S., & Iñigo-Elias, E. E. (2007). Current range of the eastern population of Painted Bunting (Passerina ciris). Part II: Winter range. North American Birds, 61 (3), 29. Retrieved from http://pubs.er.usgs.gov/publication/5224852

Sykes Jr., P. W., & Holzman, S. (2005). Current range of the eastern population of Painted Bunting (Passerina ciris). Part 1: Breeding. North American Birds, 59 (1), 14. Retrieved from http://pubs.er.usgs.gov/publication/5224505

Sykes Jr., P. W., Manfredi, L., & Padura, M. (2006). A brief report on the illegal cage-bird trade in southern Florida: A potentially serious negative impact on the eastern population of Painted Bunting (Passerina ciris). North American Birds, 60 (2), 4. Retrieved from http://pubs.er.usgs.gov/publication/5224757

