## Supplementary material for "Integrating tracking and resight data from breeding Painted Bunting populations enables unbiased inferences about migratory connectivity and winter range survival": Code for simulations, case study, and figures

### Appendix S1

Code for simulations, case study, and figures

Below, we provide all code necessary to recreate the identifiability and Painted Bunting analyses presented in the paper.

First, load necessary packages:

```
library(abind)
library(dplyr)
library(jagsUI)
library(MigConnectivity)
library(tibble)
library(purrr)
library(tidyr)
library(broom)
library(ggplot2)
```

### Identifiability analysis

#### Helper functions

To make it easier to simulate data sets under each of the scenarios presented in the paper, we first created several helper functions. The `sim_psi()` function generates random transition probabilities between a user-defined number of breeding sites and non-breeding regions:

```
sim_psi <- function(nBrSites, nWiSites){
  psi <- matrix(NA, nrow = nBrSites, ncol = nWiSites)
  for(i in 1:nBrSites){
    x <- runif(nWiSites, min = 0.2, max = 0.9)
    psi[i,] <- x/sum(x)
  }

  return(psi)
}
```

The `sim_marray()` function simulates capture histories summarized using the m-array format. User must provide the number of individuals banded during each occasion, number of years, survival probability, and recapture/resight probability:

```
sim_marray <- function(nInd, nYears, phi, d){
  if(length(d)==1){
    d <- rep(d, nYears - 1)
  }

  if(length(phi)==1){
    phi <- rep(phi, nYears - 1)
  }
```

```
  q <- 1 - d

  phi.mat <- m <- matrix(0, nrow = nYears - 1, ncol = nYears)


  for(t in 1:(nYears - 1)){
    phi.mat[t, t] <- phi[t] * d[t]

    for(j in (t + 1):nYears){
      phi.mat[t, j] <- prod(phi[t:j]) * prod(q[t:(j - 1)]) * d[j]
    }
  }

  phi.mat[,ncol(phi.mat)] <- 1 - apply(phi.mat[,-ncol(phi.mat)], 1, sum)

  ## Fill in capture histories
  m[1,] <- rmultinom(n = 1, size = nInd, prob = phi.mat[1,])

  for(t in 2:(nYears-1)){
      m[t,] <- rmultinom(n = 1, size = nInd + sum(m[, t - 1]), prob = phi.mat[t,])
  }

  return(m)
}
```

The `sim_data()` function uses the previous two functions to simulate all of the data necessary to fit the integrated tag recovery/survival model. Users can specify number of breeding sites and non-breeding regions (default is 3), number of years (default is 5), number of archival tags deployed each year (default is 30), number of individuals added to the mark-recapture data set each year (default is 100), recovery probability for archival tags (default is 0.8), and recapture/resight probability for individuals in the capture-recapture data (default is 0.6). Users may also specify the transition probabilities and non-breeding survival probabilities. If these values are not provided, random values will be generated. The function returns a named list with all data objects needed to fit the integrated model:

```
sim_data <- function(nBrSites = 3, nWiSites = 3, nYears = 5,
                     nTag = 30, nMark = 100, p = 0.8, d = 0.6,
                     psi = NULL, phi = NULL){

  ### Simulate transition matrix and winter region survivals
  if(is.null(psi))  {psi <- sim_psi(nBrSites, nWiSites)}

  if(is.null(phi)) {phi <- runif(nWiSites, min = 0.2, max = 0.9)}

    ### Weighted average apparent survival
    PHI <- apply(psi, 1, function(x) sum(x * phi))

    ### Simulate archival tag recoveries for each site x year
    ## Year 1
    W <- t(apply(psi, 1, function(x) rmultinom(1, nTag, x)))
    w <- t(apply(W, 1, function(x) rbinom(nWiSites, size = x, prob = phi * p)))
    n <- rowSums(w)

    ## Years 2:nYears
    for(t in 2:nYears){
```

```
      W2 <- t(apply(psi, 1, function(x) rmultinom(1, nTag, x)))
      w2 <- t(apply(W2, 1, function(x) rbinom(nWiSites, size = x, prob = phi*p)))
      n2 <- rowSums(w2)
      W <- abind(W, W2, along = 3)
      w <- abind(w, w2, along = 3)
      n <- cbind(n, n2)
    }

    ### Simulate CMR data
    marr <- sim_marray(nInd = nMark, nYears = nYears, phi = PHI[1], d = d)
    r <- rowSums(marr)

    for(i in 2:nBrSites){
      marr2 <- sim_marray(nInd = nMark, nYears = nYears, phi = PHI[i], d = d)
      r2 <- rowSums(marr2)
      marr <- abind(marr, marr2, along = 3)
      r <- rbind(r, r2)
    }

    ### Convert number of tags deployed each year into nBrSites x nYears matrix
    if(length(nTag)==1){
      nTag = matrix(nTag, nrow = nBrSites, ncol = nYears)
    }

    if(is.null(dim(nTag))){
      nTag = matrix(rep(nTag, nYears), ncol = nYears)
    }


    dat <- list(w = w, W.init = W, n = n, marr = marr, r = r, nTag = nTag,
                nBrSites = nBrSites, nWiSites = nWiSites, nYears = nYears,
                psi.sim = psi, phi.sim = phi, PHI.sim = PHI)

  return(dat)
}
```

#### JAGS model

The following code will create a file with the JAGS code for the integrated model:

```
sink(file="survivor_bias.jags")
cat("
    model {
    ######################## LIKELIHOOD #########################

    for(i in 1:nBrSites){
      for (t in 1:nYears){

      ### Tag recovery model
      W[i,1:nWiSites, t] ~ dmulti(psi[i,1:nWiSites], nTag[i, t])

        for(j in 1:nWiSites){
          w[i,j,t] ~ dbinom(phi[j] * p, W[i,j,t])
        }
```

```
    } # End t loop

    ### Weighted survival
    PHI[i] <- sum(psi[i, 1:nWiSites] * phi[1:nWiSites])

    ### CJS model (m-array format)
    for (t in 1:(nYears - 1)){
      marr[t, 1:nYears, i] ~ dmulti(pr[i, t, ], r[i, t])
    }

    # Define the cell probabilities of the m-array
    # Main diagonal
    for (t in 1:(nYears - 1)){
      pr[i, t, t]  <- PHI[i] * d

    # Above main diagonal
      for (j in (t + 1):(nYears - 1)){
        pr[i, t, j] <- PHI[i]^(j - t + 1) * q^(j-t) * d
      } #j

    # Below main diagonal
      for (j in 1:(t - 1)){
        pr[i, t, j] <- 0
      } #j
    } #t

    # Last column: probability of non-recapture
    for (t in 1:(nYears - 1)){
      pr[i, t, nYears] <- 1 - sum(pr[i, t, 1:(nYears - 1)])
    } #t
  } # End i loop

  ####################### PRIORS ###############################
    ### Tag recovery probability
    p ~ dunif(0, 1)

    ### Resight probability
    d ~ dunif(0, 1)
    q <- 1 - d

  for(i in 1:nBrSites){
    for(j in 1:nWiSites){
      psi_raw[i, j] ~ dgamma(1, 1)
      psi[i, j] <- psi_raw[i, j] / sum(psi_raw[i, 1:nWiSites])
    }
  }

  for(j in 1:nWiSites){
      phi[j] ~ dunif(0, 1)
  }

  } # End model
  ", fill=TRUE)
```

```
sink()
```

#### Identifiability analysis

First, we set the number of simulations to run and number of breeding sites/non-breeding regions. In order to calculate the strength of migratory connectivity, we also need to define distance between sites/regions and the relative abundance of the breeding sites:

```
## Number of simulations
nSims <- 25


### Breeding/non-breeding sites
nBreeding <- 3
nNonBreeding <- 3

breedDist <- matrix(c(0, 1, 2,
                      1, 0, 1,
                      2, 1, 0), nBreeding, nBreeding)
nonBreedDist <- matrix(c(0, 1, 2,
                         1, 0, 1,
                         2, 1, 0), nNonBreeding, nNonBreeding)

relN <- rep(1/nBreeding, nBreeding)
```

Next, we will create an empty data frame to store results from each simulation. Each element of the `bias` and `psi` columns are themselves data frames that store results for each non-breeding region:

```
identify_results <- as_data_frame(data.frame(cv = numeric(length = nSims),
                                             MC.true = numeric(length = nSims),
                                             MC.raw = numeric(length = nSims),
                                             MC.est = numeric(length = nSims)))

identify_results <- mutate(identify_results, bias = list(length = nSims),
                           psi = list(length = nSims))
```

Now set MCMC settings, including the parameters to monitor, number of chains, number iterations, length of the burn-in period:

```
## Parameters to monitor
params <- c("psi", "phi", "PHI", "p", "d")

## MCMC settings
nc <- 3       ## Number of chains
ni <- 500   ## Number of iterations
na <- 100    ## Length adaptation phase
nb <- 100    ## Length of burn-in phase
nt <- 1       ## Thinning rate
```

Finally, loop over `nSims` to generate data sets, fit the model, and estimate parmaters of interest:

```
for(j in 1:nSims){
  ## Simulate data set
  dat <- sim_data(nBrSites = nBreeding, nWiSites = nNonBreeding,
                  nTag = 10000, nMark = 20000)
```

```
  ## Estimate CV of non-breeding survivals
  identify_results$cv[j] <- sd(dat$phi)/mean(dat$phi)

  ## Calculate true MC
  identify_results$MC.true[j] <- round(MigConnectivity::calcMC(breedDist,
                                                                nonBreedDist,
                                                                relN,
                                                                psi = dat$psi.sim),  3)

  ## Estimate psi from the raw recovery data
  raw.psi <- t(apply(rowSums(dat$w, dims = 2), 1, function(x) x/sum(x)))

  ## Calculate MC from the raw psi estimate
  identify_results$MC.raw[j] <- round(MigConnectivity::calcMC(breedDist,
                                                              nonBreedDist,
                                                              relN,
                                                              psi = raw.psi), 3)

  ## Create initial values for W (necessary to prevent error message)
  inits <- function(){list(W = dat$W.init)}

  ## Fit the integrated model
  fit <- jags(data = dat, inits = inits, parameters.to.save = params,
                    model.file = "survivor_bias.jags",
                    n.chains = nc, n.adapt = na, n.iter = ni,
                    n.burnin = nb, n.thin = nt,
                    parallel = TRUE, verbose = FALSE)

  ## Caclulate MC from estimated psi
  identify_results$MC.est[j] <- round(MigConnectivity::calcMC(breedDist,
                                                              nonBreedDist,
                                                              relN,
                                                              psi = fit$mean$psi), 3)

  ## Save the true, raw, and estimated psi's
  identify_results$psi[[j]] <- data.frame(psi.true = c(dat$psi.sim),
                                          psi.raw = c(raw.psi),
                                          psi.est = c(fit$mean$psi))

  ## Calculate bias of psi estimates
  identify_results$bias[[j]] <- data.frame(phi = rep(dat$phi.sim, 2),
                                    phi.est = c(rep(NA, 3), fit$mean$phi),
                                    value = c(apply((raw.psi - dat$psi.sim)/dat$psi.sim,
                                                    2, mean),
                                              apply((fit$mean$psi - dat$psi.sim)/dat$psi.sim,
                                                    2, mean)),
                                    Estimate = rep(c("Raw", "Corrected"),
                                                   each = nNonBreeding))
  }

## Save results
saveRDS(identify_results, "identify_results.rds")
```

Using this code, users can change input values to recreate estimability scenarios from the paper or explore different scenarios.

#### Evaluating model output

In the manuscript, we present the slopes coefficients from a regression of bias vs. non-breeding survival. The following code will estimate those coefficients from the simulation above (exact values may be slightly different from those in the manuscript due to random generation of simulated data sets):

```
## Read in model output
identify_results <- readRDS("identify_results.rds")

## Pull out the bias column
identify_df <- map_df(identify_results$bias, .f = rbind)

## Fit model to all values from raw and integrated estimates
identify_slope <- identify_df %>%
  nest(-Estimate) %>%
  mutate(fit = map(data, ~ lm(value ~ phi, data = .)),
         slope = map(fit, coef)) %>%
  unnest(c(slope)) %>%
  slice(which(row_number() %% 2 == 0))

## Remove data and fit columns
identify_slope <- select(identify_slope, -data, -fit)

## Slope coefficient from raw estimates
round(identify_slope$slope[1], 2)
```

```
## [1] 1.25
```

```
## Slope coefficient from integrated model estimates
round(identify_slope$slope[2], 2)
```

```
## [1] 0.11
```

The following code will recreate Figure 1 from the paper (change `nSims` to exactly replicate analysis in the main text) showing the relationship between non-breeding survival and bias in the $\psi$ estimates from the raw and corrected data:

```
## Create new column for axis labels
identify_df <- dplyr::mutate(identify_df,
                      Estimate = case_when(Estimate == "Raw" ~ "Uncorrected",
                                           Estimate == "Corrected" ~ "Integrated model"))

## Figure code
ggplot(identify_df, aes(x = phi, y = value, color = Estimate,
                        linetype = Estimate, alpha = Estimate)) +
  geom_hline(yintercept = 0, linetype = "longdash", alpha = 0.5) +
  geom_point(size = 2.5) +
  stat_smooth(method = "lm", se = FALSE, color = "grey10", alpha = 1, size = 1.25) +
  scale_color_manual(values = c("#446E9B", "#D47500")) +
  scale_shape_manual(values = c(16, 21)) +
  scale_alpha_manual(values = c(0.7, 0.7)) +
  scale_linetype_manual(values = c("dashed", "solid")) +
  theme_classic() +
  scale_x_continuous(expression(paste("Non-breeding survival (", phi[j], ")"))) +
  scale_y_continuous("Bias", limits = c(-1, 1.4)) +
```

```r
  theme(legend.title = element_blank())
```

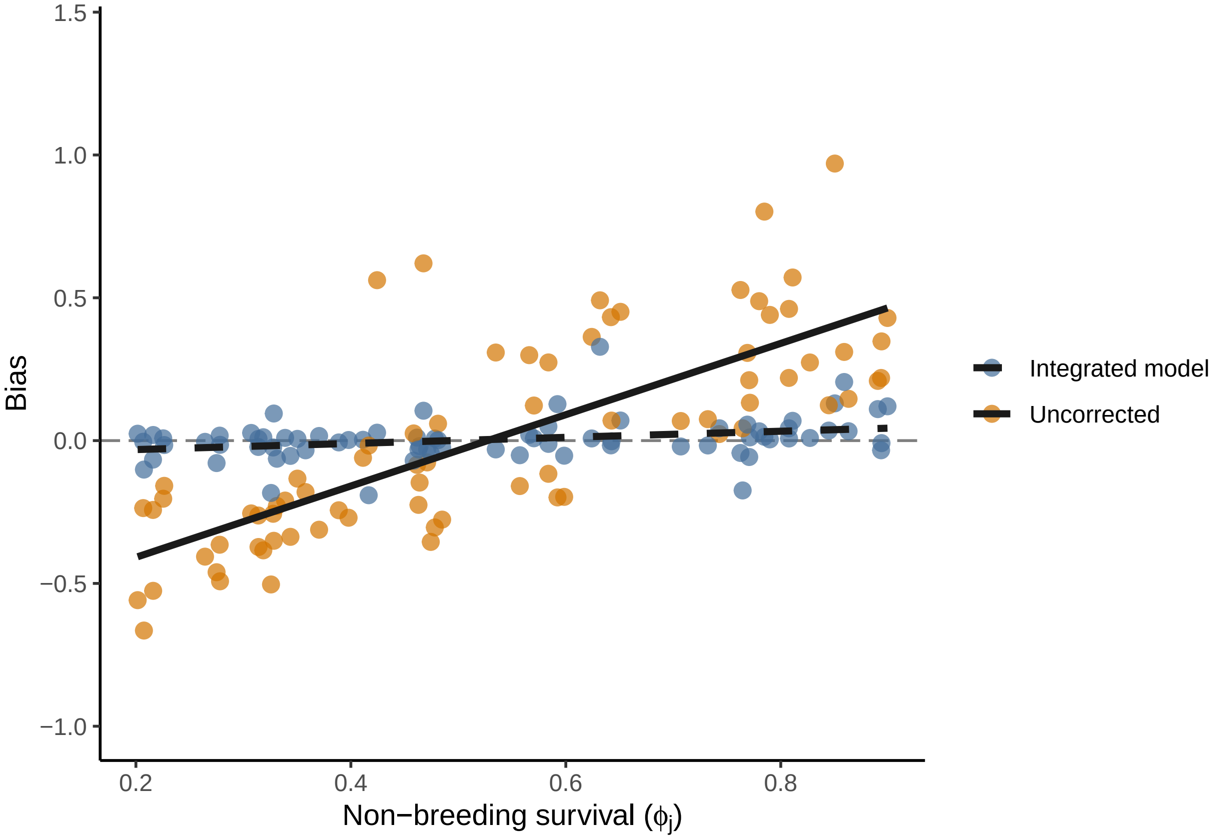


The following code will recreate Figure 2 from the paper showing the relationship between the true and estimated MC from the raw and corrected $\psi$ estimates:

```r
ggplot(identify_results) +
  geom_abline(intercept = 0, slope = 1) +
  geom_point(aes(x = MC.true, y = MC.raw, color = "#D47500"), size = 3, alpha = 0.7) +
  geom_point(aes(x = MC.true, y = MC.est, color = "#446E9B"), size = 3, alpha = 0.7) +
  theme_classic() +
  scale_x_continuous("True MC") +
  scale_y_continuous("Estimated MC") +
  scale_color_identity(guide = "legend",
                       breaks = c("#D47500", "#446E9B"),
                       labels = c( "Uncorrected", "Integrated model")) +
  theme(legend.title = element_blank())
```

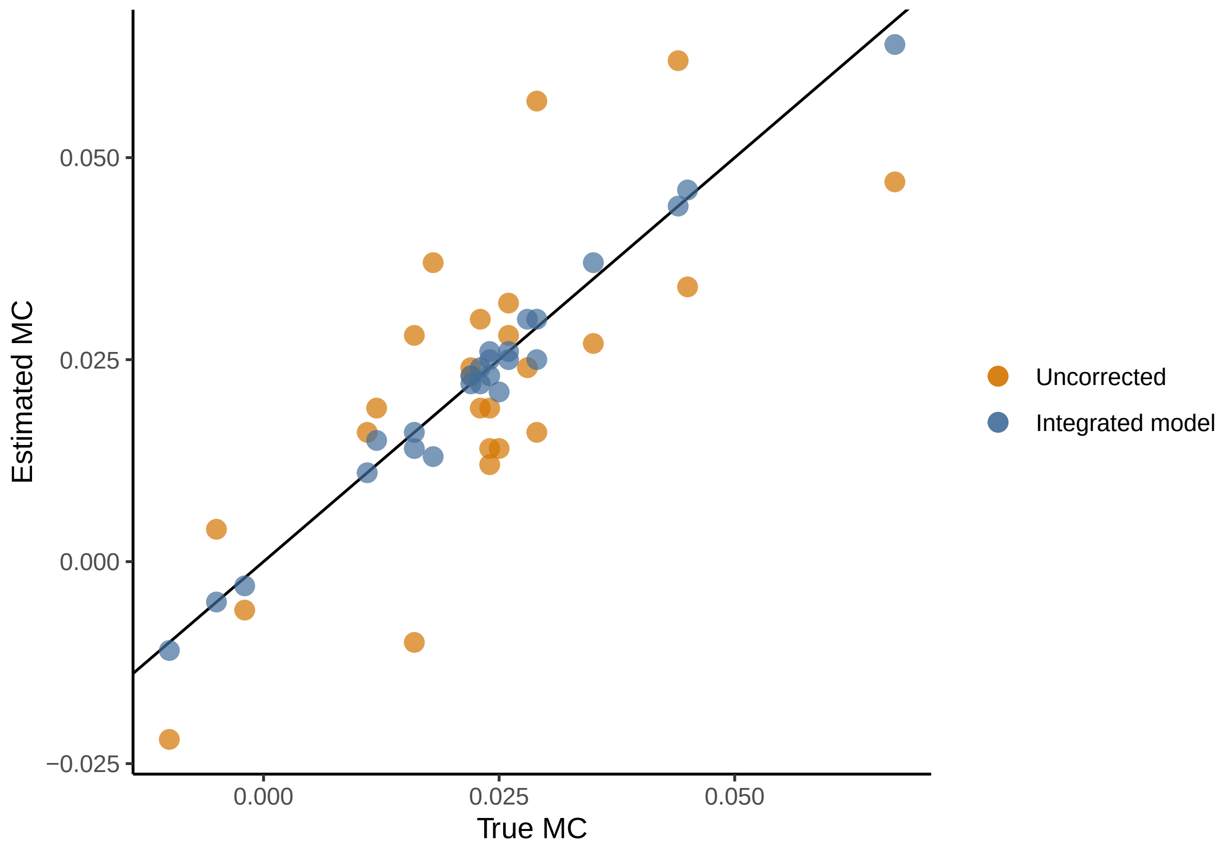
Estimated MC
0.050
0.025
0.000
−0.025
0.000
0.025
0.050
True MC
Uncorrected
Integrated model

### Case study: Migratory connectivity and

#### Geolocator data

##### Aimee add summary text

Table S1 1: Summary of geolocator deployment and recovery data. Sample sizes (n) for each site indicate the total number of light-level geolocators deployed on adult male Painted Buntings at each site. Values in each cell are the observed number of individuals from each deployment site that wintered in each non-breeding region.

| | Florida (n = 45) | Georgia (n = 41) | South Carolina (n = 67) | North Carolina (n = 30) |
|---|---|---|---|---|
| North Florida | 4 | 4 | 6 | 0 |
| South Florida | 3 | 7 | 8 | 0 |
| Cuba | 4 | 3 | 5 | 4 |
| Bahamas | 5 | 5 | 2 | 1 |

#### Mark-resight data

Table S1 2: Locations of study sites included in the capture-resight analysis. See Sykes et al. (2019) for additional details about site selection and sampling design

| Site | State | Latitude | Longitude |
|---|---|---|---|
| Ft. Caroline | FL | 30.380 | -81.482 |
| Ft. George Island | FL | 30.423 | -81.440 |
| Little Talbot Island State Park | FL | 30.456 | -81.415 |
| Cedar Point | FL | 30.441 | -81.477 |
| Blackbeard NWR | GA | 31.515 | -81.196 |
| Harris Neck NWR | GA | 31.630 | -81.269 |
| St. Catherines Is. | GA | 31.627 | -81.152 |
| ACE Basin NWR | SC | 32.665 | -80.393 |
| James Island | SC | 32.724 | -79.955 |
| Bald Head Island | NC | 33.857 | -77.985 |
| Carolina Beach State Park | NC | 34.046 | -77.910 |

#### JAGS model

```
sink(file = "pabu_model_weighted.jags")
cat("
    model {
    ######################## LIKELIHOOD #########################

    ### Survival model
    for (i in 1:nind){
      # Define latent state at first capture
      z[i, f[i]] <- 1

      for (t in (f[i]+1):n.occasions){
        # State process
        z[i, t] ~ dbern(mu1[i, t])
        mu1[i, t] <- phi[i, t-1] * z[i, t-1]
```

```
    # Observation process
    y[i, t] ~ dbern(mu2[i, t])
    mu2[i, t] <- p[i, t-1] * z[i, t]
  } #i
} #t

### Tag recovery model
  for(s in 1:nBrSites){
    # State process
      W[s, 1:nWiSites] ~ dmulti(psi[s, 1:nWiSites], nTag_total[s])

  for(j in 1:nWiSites){
    # Observation process
      w[s, j] ~ dbinom(phi.w[j] * d, W[s, j])
    } #j
  } #s

### Posterior predictive check
  for (i in 1:nind){
    for (t in (f[i] + 1):n.occasions){
    #simulated obs for as many individuals as in data set, each date
    y_sim[i, t]~ dbern(mu2[i, t])
  } #t

  ind.obs[i] <- sum(y[i, f[i]:n.occasions]) ### total number of years ind[i] seen
  ind.obs_sim[i] <- sum(y_sim[i, f[i]:n.occasions])
  ind.obs_exp[i] <- sum(mu2[i, (f[i]+1):n.occasions]) + 1 ### expected total
  depobs[i] <- pow((pow(ind.obs[i], 0.5) - pow(ind.obs_exp[i], 0.5)), 2)

  # Freeman-Tukey measure of departure from expected, observed data
  # departure from expected,
  depsim[i] <- pow((pow(ind.obs_sim[i], 0.5) - pow(ind.obs_exp[i], 0.5)), 2)
} #i

PABUfit <- sum(depobs[]) #discrepancy, observed data
PABUfit.sim <- sum(depsim[]) #discrepancy, simulated data

####################### PRIORS AND CONSTRAINTS ###############################

### Non-breeding region survival
for(j in 1:nWiSites){
  phi.w[j] ~ dunif(0, 1)
} #j

### Breeding region survival (weighted)
for(s in 1:nBrSites){
    phi.r[s] <- sum(psi[s ,1:nWiSites] * phi.w[1:nWiSites])  # Weighted survival
    lphi.r[s] <- log(phi.r[s] / (1 - phi.r[s]))  # Logit survival
} #s

### Feeder survival with development effect
for(i in 1:nFeeder){
  epsilon.f[i] ~ dnorm(0, tau.f)
```

```
    # beta.700 created in model, dev.700 in as data
    lphi.f[i] <- lphi.r[region[i]] +  beta.700 * dev.700[i] + epsilon.f[i]

    # mean survival for mean latitude, mean development
    phi.f[i] <- 1/(1 + exp(-lphi.f[i]))
  } #i

  sigma.f ~ dunif(0, 10)
  tau.f <- pow(sigma.f, -2)
  sigma2.f <- pow(sigma.f, 2)

  ### Slope coefficient; effect of development on feeder-specific survival
  beta.700 ~ dnorm(0, 0.37)I(-10, 10)

  ### Detection probability for marked individuals
  for (i in 1:nind){
    for (t in f[i]:(n.occasions-1)){
      phi[i, t] <- phi.f[feeder[i]]
      logit(p[i, t]) <- gamma.p + gamma.p.bo*banding_only[i, t] +
                        gamma.p.xtra*extra_hrs[i, t] +
                        gamma.p.nogo*nogo[i, t] + epsilon.p[i]
    } #t
  } #i

  ### Random effect of individual on detection
  for (i in 1:nind){
    epsilon.p[i] ~ dnorm(0, tau.p)
  } #i

  sigma.p ~ dunif(0,10)
  tau.p <- pow(sigma.p, -2)
  sigma2.p <- pow(sigma.p, 2)

  ### For recapture parameters
  gamma.p ~ dnorm(0, 0.37)I(-10, 10)
  gamma.p.bo ~ dnorm(0, 0.37)I(-10, 10)
  gamma.p.xtra ~ dnorm(0, 0.37)I(-10, 10)
  gamma.p.nogo ~ dnorm(0, 0.37)I(-10, 10)
  p.mean <- 1/(1 + exp(-gamma.p))
  p.xtra <- 1/(1 + exp(-gamma.p - gamma.p.xtra))
  p.nogo <- 1/(1 + exp(-gamma.p - gamma.p.nogo))
  p.bo <- 1/(1 + exp(-gamma.p - gamma.p.bo))

  ### Tag recovery probability (assume constant)
  d ~ dunif(0, 1)

  ### Transition probability between regions
  for(s in 1:nBrSites){
    for(j in 1:nWiSites){
      psi_raw[s, j] ~ dgamma(1, 1)
      psi[s, j] <- psi_raw[s, j] / sum(psi_raw[s, 1:nWiSites])
    } #j
  } #s
```

```
    ### Set up post-predictive check
    for (i in 1:nind){
      y_sim[i, f[i]] <- 1
    } #i

    }
    ",fill = TRUE)
sink()
```

#### Model fitting

#### Maps

The following code will recreate Figure 4 from the paper:

```
library(ggforce)
library(ggplot2)
library("rnaturalearth")
library("rnaturalearthdata")
library("ggspatial")

target_crs <- "+proj=longlat +datum=WGS84 +no_defs +ellps=WGS84 +towgs84=0,0,0"

deployment_xy <- sf::st_sfc(sf::st_point(c(-77.9083, 34.0167)),
                            sf::st_point(c(-80.0817, 32.6086)),
                            sf::st_point(c(-80.8264, 32.3293)),
                            sf::st_point(c(-81.2417, 31.4775)),
                            sf::st_point(c(-81.4153, 30.4591)),
                            crs = target_crs)

deployment_df <- data.frame(site = c("NC", "SC1", "SC2", "GA", "FL"))
deployment_sf <- sf::st_sf(deployment_df, geometry = deployment_xy)

cmr_xy <- sf::st_sfc(sf::st_point(c(-81.482202, 30.380148)),
                     sf::st_point(c(-81.440610, 30.423640)),
                     sf::st_point(c(-81.415803, 30.456306)),
                     sf::st_point(c(-81.477359, 30.441586)),
                     sf::st_point(c(-81.196400, 31.515233)),
                     sf::st_point(c(-81.269455, 31.630575)),
                     sf::st_point(c(-81.152426, 31.626685)),
                     sf::st_point(c(-80.393065, 32.664867)),
                     sf::st_point(c(-79.955873, 32.724422)),
                     sf::st_point(c(-77.985856, 33.857130)),
                     sf::st_point(c(-77.910856, 34.045841)),
                     crs = target_crs)


cmr_df <- data.frame(State = c("FL", "FL", "FL", "FL",
                               "GA", "GA", "GA",
                               "SC", "SC",
                               "NC", "NC"),
                     Site = c("Ft. Caroline", "Ft. George Island",
                              "Little Talbot Island State Park",
                              "Cedar Point", "Blackbeard NWR",
```

```r
                              "Harris Neck NWR", "St. Catherines Is.",
                              "ACE Basin NWR", "James Island",
                              "Bald Head Island", "Carolina Beach State Park"))

cmr_sf <- sf::st_sf(cmr_df, geometry = cmr_xy)


world <- ne_countries(scale = "medium", returnclass = "sf")
us <- USAboundaries::us_states()

cuba <- dplyr::filter(world, name == "Cuba")
bahamas <- dplyr::filter(world, name == "Bahamas")
sfl <- dplyr::filter(us, name == "Florida")
sfl <- sf::st_crop(sfl, ymin = 18, ymax = 28.5, xmin = -86, xmax = -71)
nfl <- dplyr::filter(us, name == "Florida" | name == "Georgia")
nfl <- sf::st_union(nfl, by_feature = TRUE)
nfl <- sf::st_crop(nfl, ymin = 28.5, ymax = 31, xmin = -83, xmax = -71)



################
### Base map
################
worldmap_trans <- sf::st_transform(world, crs = target_crs)

disp_win_wgs84 <- sf::st_sfc(sf::st_point(c(-85, 19.5)), sf::st_point(c(-72, 34)),
                         crs = 4326)

disp_win_trans <- sf::st_transform(disp_win_wgs84, crs = target_crs)
disp_win_coord <- sf::st_coordinates(disp_win_trans)

base <- ggplot() +
  theme_bw() +
  geom_sf(data = worldmap_trans, color = NA, fill = "grey70", size = 0.4) +
  geom_sf(data = us, color = "grey80", fill = "grey70", size = 0.4) +
  geom_sf(data = cuba, fill = "#D9AF6B", color = "grey20") +
  geom_sf(data = bahamas, fill = "#9C9C5E", color = "grey20") +
  geom_sf(data = nfl, fill = "#467378", color = "grey20") +
  geom_sf(data = sfl, fill = "#AF6458", color = "grey20") +
  coord_sf(xlim = disp_win_coord[,'X'], ylim = disp_win_coord[,'Y'],
           datum = target_crs) +
  scale_y_continuous(breaks = c(18, 22, 26, 30, 34)) +
  theme(panel.background = element_rect(fill = "grey98")) +#c8d7d4
  scale_color_manual(values = c("#9C9C5E", "#D9AF6B", "#467378","#AF6458")) +
  guides(color = FALSE) +
  scale_size(range = c(0.75, 3))


################
### NC map
################
blank_df <- data.frame(psi.range = c(0.1, 0.59))
```

```r
NCbeziers <- data.frame(
  x = c(-77.9083, -75, -77,
        -77.9083, -72, -77,
        -77.9083, -78, -80.8,
        -77.9083, -76, -79.8),
  y = c(34.0167, 30, 27,
        34.0167, 26.75, 22,
        34.0167, 30, 29.5,
        34.0167, 28, 26.75),
  target = rep(c("Bahamas", "Cuba", "North Florida", "South Florida"), each = 3),
  psi = rep(c(0.20, 0.59, 0.11, 0.10), each = 3)
)


nc <- base +
  geom_bezier(aes(x = x, y = y, group = target, color = target, size = psi),
              arrow = arrow(length = unit(c(0.4, 0.7, 0.3, 0.3), "cm")),
              data = NCbeziers) +
  geom_sf(data = deployment_sf[deployment_sf$site == "NC",],
          shape = 21, fill = "#ca562c", size = 3) +
  geom_sf(data = cmr_sf[cmr_sf$State == "NC",], color = "grey10", size = 3, shape = 4) +
  theme(axis.title = element_blank(), axis.text = element_blank()) +
  geom_text(data = NCbeziers[3,], aes(x = x, y = y, label = psi, color = target),
            nudge_x = 0.85, nudge_y = -0.275, size = 5, fontface = "bold") +
  geom_text(data = NCbeziers[6,], aes(x = x, y = y, label = psi, color = target),
            nudge_x = 1, nudge_y = -0.35, size = 5, fontface = "bold") +
  geom_text(data = NCbeziers[9,], aes(x = x, y = y, label = psi, color = target),
            nudge_x = 0.75, nudge_y = -0.4, size = 5, fontface = "bold") +
  geom_text(data = NCbeziers[12,], aes(x = x, y = y, label = psi, color = target),
            nudge_x = 0.55, nudge_y = -0.5, size = 5, fontface = "bold") +
  guides(size = FALSE) +
  annotation_north_arrow(location = "tr", which_north = "true",
                         pad_x = unit(0.1, "in"), pad_y = unit(0.1, "in"),
                         style = north_arrow_fancy_orienteering) +
  coord_sf(xlim = disp_win_coord[,'X'], ylim = disp_win_coord[,'Y'],
           datum = target_crs) +
  geom_blank(data = blank_df, aes(size = psi.range))

################
### SC map
################

target_df <- data.frame(x = c(-84.25, -84, -80.75, -73.2),
                        y = c(28.6, 26.41, 20.4, 23.4),
                        phi = c("phi == \n 0.73", "phi == \n 0.76",
                                "phi == 0.57", "phi == 0.72"),
                        site = c("North Florida", "South Florida",
                                 "Cuba", "Bahamas"))

SCbeziers <- data.frame(
  x = c(-80.0817, -76, -77,
        -80.0817, -72, -77,
        -80.0817, -79, -80.8,
```

```
        -80.0817, -76, -79.7),
  y = c(32.6086, 30.5, 26.75,
        32.6086, 30, 22,
        32.6086, 30.5, 29.5,
        32.6086, 29, 26),
  target = rep(c("Bahamas", "Cuba", "North Florida", "South Florida"), each = 3),
  psi = rep(c(0.12, 0.28, 0.27, 0.32), each = 3)
)


sc <- base +
  geom_bezier(aes(x = x, y = y, group = target, color = target, size = psi),
              arrow = arrow(length = unit(c(0.25, 0.5, 0.4, 0.45), "cm"), angle = 40),
              data = SCbeziers) +
  geom_sf(data = deployment_sf[deployment_sf$site %in% c("SC1", "SC2"),],
          shape = 21, fill = "#ca562c", size = 3) +
  geom_sf(data = cmr_sf[cmr_sf$State == "SC",], color = "grey10", size = 3, shape = 4) +
  theme(axis.title = element_blank(),
        axis.text.x = element_blank()) +
  geom_text(data = target_df, aes(x = x, y = y, label = phi),
            color = "grey10", fontface = "bold", size = 5.5, parse = TRUE) +
  geom_segment(aes(x = -83.1, xend = -82, y = 28.7, yend = 29.5), color = "grey20") +
  geom_segment(aes(x = -82.7, xend = -81.2, y = 26.41, yend = 27), color = "grey20") +
  geom_segment(aes(x = -80.3, xend = -80, y = 20.8, yend = 22), color = "grey20") +
  geom_text(data = SCbeziers[3,], aes(x = x, y = y, label = psi, color = target),
            nudge_x = 0.85, nudge_y = -0.275, size = 5, fontface = "bold") +
  geom_text(data = SCbeziers[6,], aes(x = x, y = y, label = psi, color = target),
            nudge_x = 1, nudge_y = -0.3, size = 5, fontface = "bold") +
  geom_text(data = SCbeziers[9,], aes(x = x, y = y, label = psi, color = target),
            nudge_x = 0.75, nudge_y = -0.4, size = 5, fontface = "bold") +
  geom_text(data = SCbeziers[12,], aes(x = x, y = y, label = psi, color = target),
            nudge_x = 0.55, nudge_y = -0.5, size = 5, fontface = "bold") +
  guides(size = FALSE) +coord_sf(xlim = disp_win_coord[,'X'], ylim = disp_win_coord[,'Y'],
                                 datum = target_crs) +
  geom_blank(data = blank_df, aes(size = psi.range))


################
### GA map
################

GAbeziers <- data.frame(
  x = c(-81.2417, -77, -77.5,
        -81.2417, -73, -77,
        -81.2417, -79.75, -80.8,
        -81.2417, -77, -79.7),
  y = c(31.4775, 30.5, 27.25,
        31.4775, 30, 22,
        31.4775, 30.75, 29.5,
        31.4775, 29, 26),
  target = rep(c("Bahamas", "Cuba", "North Florida", "South Florida"), each = 3),
  psi = rep(c(0.25, 0.21, 0.21, 0.32), each = 3)
)
```

```
ga <- base +
  geom_bezier(aes(x = x, y = y, group = target, color = target, size = psi),
              arrow = arrow(length = unit(c(0.45, 0.4, 0.4, 0.5), "cm"), angle = 40),
              data = GAbeziers) +
  geom_sf(data = deployment_sf[deployment_sf$site == "GA",],
          shape = 21, fill = "#ca562c", size = 3) +
  geom_sf(data = cmr_sf[cmr_sf$State == "GA",], color = "grey10", size = 3, shape = 4) +
  theme(axis.title = element_blank()) +
  geom_text(data = GAbeziers[3,], aes(x = x, y = y, label = psi, color = target),
            nudge_x = 0.85, nudge_y = -0.275, size = 5, fontface = "bold") +
  geom_text(data = GAbeziers[6,], aes(x = x, y = y, label = psi, color = target),
            nudge_x = 0.7, nudge_y = -0.275, size = 5, fontface = "bold") +
  geom_text(data = GAbeziers[9,], aes(x = x, y = y, label = psi, color = target),
            nudge_x = 0.75, nudge_y = -0.4, size = 5, fontface = "bold") +
  geom_text(data = GAbeziers[12,], aes(x = x, y = y, label = psi, color = target),
            nudge_x = 0.55, nudge_y = -0.5, size = 5, fontface = "bold") +
  annotation_scale(location = "bl", width_hint = 0.5) +
  guides(size = FALSE) + coord_sf(xlim = disp_win_coord[,'X'],
                                  ylim = disp_win_coord[,'Y'],
                                  datum = target_crs) +
  geom_blank(data = blank_df, aes(size = psi.range))

################
### FL map
################

FLbeziers <- data.frame(
  x = c(-81.4153, -77.5, -77.5,
        -81.4153, -73, -77,
        -81.4153, -79.9, -80.7,
        -81.4153, -78, -79.8),
  y = c(30.4591, 30.5, 27.25,
        30.4591, 30, 22,
        30.4591, 30.1, 29,
        30.4591, 30, 26.75),
  target = rep(c("Bahamas", "Cuba", "North Florida", "South Florida"), each = 3),
  psi = rep(c(0.29, 0.27, 0.25, 0.20), each = 3)
)


fl <- base +
  geom_bezier(aes(x = x, y = y, group = target, color = target, size = psi),
              arrow = arrow(length = unit(c(0.45, 0.45, 0.45, 0.35), "cm"), angle = 40),
              data = FLbeziers) +
  geom_sf(data = deployment_sf[deployment_sf$site == "FL",],
          shape = 21, fill = "#ca562c", size = 3) +
  geom_sf(data = cmr_sf[cmr_sf$State == "FL",],
          color = "grey10", size = 3, shape = 4) +
  theme(axis.title = element_blank(),
        axis.text.y = element_blank(),
        legend.position = c(0.875, 0.75),
        legend.background = element_rect(fill = NA, color = "grey20"),
```

```
        legend.key  = element_rect(fill = NA),
        legend.title = element_text(hjust = 0.5)) +
  guides(size = "none") +
  geom_text(data = FLbeziers[3,], aes(x = x, y = y, label = psi, color = target),
            nudge_x = 0.8, nudge_y = -0.275, size = 5, fontface = "bold") +
  geom_text(data = FLbeziers[6,], aes(x = x, y = y, label = psi, color = target),
            nudge_x = 1, nudge_y = -0.3, size = 5, fontface = "bold") +
  geom_text(data = FLbeziers[9,], aes(x = x, y = y, label = psi, color = target),
            nudge_x = 0.75, nudge_y = -0.2, size = 5, fontface = "bold") +
  geom_text(data = FLbeziers[12,], aes(x = x, y = y, label = psi, color = target),
            nudge_x = 0.6, nudge_y = -0.6, size = 5, fontface = "bold") +
  coord_sf(xlim = disp_win_coord[,'X'], ylim = disp_win_coord[,'Y'], datum = target_crs) +
  geom_blank(data = blank_df, aes(size = psi.range))


psi_maps <- cowplot::plot_grid(sc, nc, ga, fl,
                               nrow = 2, ncol = 2, align = 'hv',
                               labels = "AUTO")
psi_maps
```
