## Supplementary tables and figures for "Integrating tracking and resight data from breeding Painted Bunting populations enables unbiased inferences about migratory connectivity and winter range survival"

### Appendix S2

#### Supplementary tables and figures

Table S2 1: Estimability of transition probabilities and non-breeding survival as a function of the number of years in the capture-mark-recapture study. Correlations are the mean and 95% confidence interval from the 500 simulated data sets

| Number of years | Correlation - integrated model | Correlation - uncorrected | Bias in $\phi_j$ |
|---|---|---|---|
| 4 | 0.449 (0.401 - 0.497) | 1.249 (1.192 - 1.307) | 0.0411769 |
| 6 | 0.373 (0.329 - 0.418) | 1.244 (1.188 - 1.299) | 0.0454857 |
| 8 | 0.34 (0.296 - 0.384) | 1.272 (1.217 - 1.326) | 0.0399037 |
| 10 | 0.286 (0.247 - 0.324) | 1.248 (1.195 - 1.301) | 0.0305845 |
| 12 | 0.309 (0.269 - 0.349) | 1.254 (1.202 - 1.306) | 0.0298491 |

Figure S2 1: Correlation between non-breeding survival and bias in the estimated transition probabilities as a function of the number of years

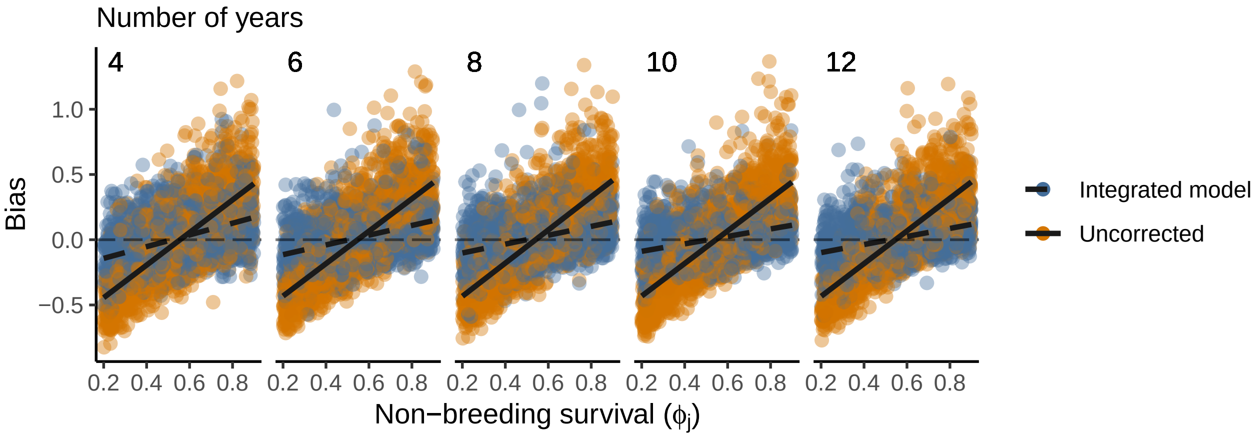

Table S2 2: Estimability of transition probabilities and non-breeding survival as a function of the number of breeding sites and non-breeding regions. Correlations are the mean and 95% confidence interval from the 500 simulated data sets

| Number of Sites | Correlation - integrated model | Correlation - uncorrected | Bias in $\phi_j$ |
|---|---|---|---|
| 3 | 0.438 (0.392 - 0.483) | 1.265 (1.21 - 1.32) | 0.0429147 |
| 4 | 0.447 (0.409 - 0.486) | 1.409 (1.364 - 1.454) | 0.0414064 |
| 5 | 0.504 (0.472 - 0.537) | 1.503 (1.464 - 1.542) | 0.0371564 |
| 6 | 0.524 (0.497 - 0.552) | 1.576 (1.542 - 1.61) | 0.0371855 |
| 7 | 0.477 (0.451 - 0.502) | 1.572 (1.54 - 1.604) | 0.0381506 |

Figure S2 2: Correlation between non-breeding survival and bias in the estimated transition probabilities as a function of the number of sites

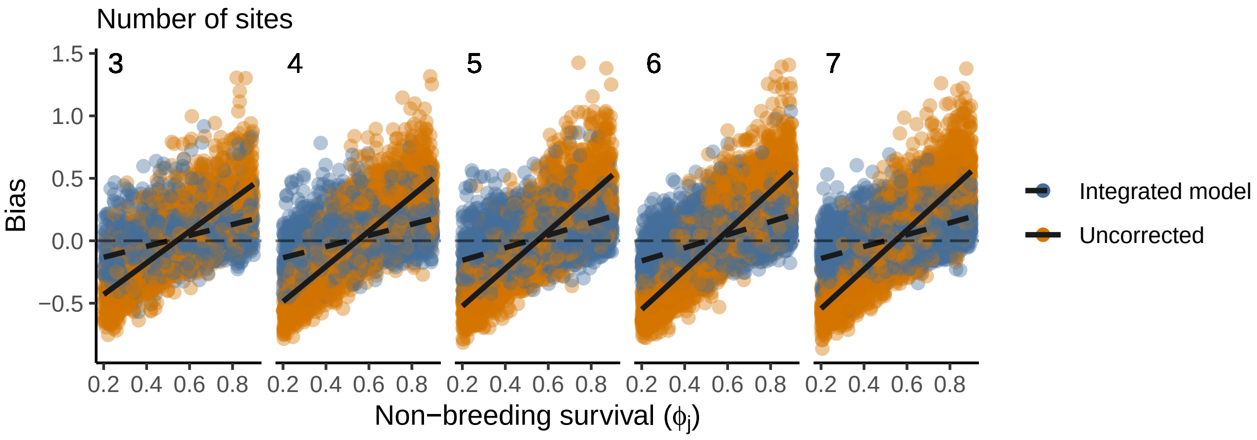
